# Solid tumor growth depends on an intricate equilibrium of malignant cell states

**DOI:** 10.1101/2023.12.30.573100

**Authors:** Stefan R. Torborg, Olivera Grbovic-Huezo, Anupriya Singhal, Matilda Holm, Katherine Wu, Xuexiang Han, Yu-Jui Ho, Caj Haglund, Michael J. Mitchell, Scott W. Lowe, Lukas E. Dow, Kenneth L. Pitter, Francisco J. Sanchez-Rivera, Andre Levchenko, Tuomas Tammela

## Abstract

Control of cell identity and number is central to tissue function, yet principles governing organization of malignant cells in tumor tissues remain poorly understood. Using mathematical modeling and candidate-based analysis, we discover primary and metastatic pancreatic ductal adenocarcinoma (PDAC) organize in a stereotypic pattern whereby PDAC cells responding to WNT signals (WNT-R) neighbor WNT-secreting cancer cells (WNT-S). Leveraging lineage-tracing, we reveal the WNT-R state is transient and gives rise to the WNT-S state that is highly stable and committed to organizing malignant tissue. We further show that a subset of WNT-S cells expressing the Notch ligand DLL1 form a functional niche for WNT-R cells. Genetic inactivation of WNT secretion or Notch pathway components, or cytoablation of the WNT-S state disrupts PDAC tissue organization, suppressing tumor growth and metastasis. This work indicates PDAC growth depends on an intricately controlled equilibrium of functionally distinct cancer cell states, uncovering a fundamental principle governing solid tumor growth and revealing new opportunities for therapeutic intervention.

## Introduction

Tissues are composed of distinct cellular differentiation states maintained at constant relative numbers and densities, which is essential to tissue function. For example, acinus-duct units in the pancreas are composed of acinar and ductal cells, whereas intestinal crypt-villus units are comprised of multiple specialized epithelial cell types that arise from adult stem cells (ASCs) residing in the crypt bottom^1,2^. The identity and proportions of different cell types within these tissues is carefully controlled by cell-cell interactions and other microenvironmental signaling cues. This coordinated organization facilitates cooperation between distinct cell types, enabling the various functions of healthy organs. Epithelial tissues harbor remarkable capacity for restoring cellular organization and density upon injury via activation of regenerative programs^1–3^. Whether similar organization and control of cell density takes place in tumors remains unknown.

Organogenesis during development is orchestrated by intricately controlled spatiotemporal interactions between cells, whereby precursor cells give rise to differentiated cells in carefully controlled numbers. The differentiated cells assemble into functional tissue patterns, which are stereotypic in density and cellular composition. In adult tissues, homeostasis is maintained by progenitors or ASCs, which self-renew and proliferate to sustain tissue architecture, as dictated by the demands of cell turnover within the tissue^1,3,4^. In contrast to organogenesis and tissue regeneration, our understanding of the cell-cell interactions and other cues controlling differentiation of cancer cells remains at an early stage.

Tumors are composed of functionally and molecularly distinct cancer cell differentiation states. The clinical relevance of this heterogeneity manifests in the distinct capacities of cancer cell states for growth, differentiation, treatment resistance, and metastasis. The emergence of single-cell genomics has enabled the unsupervised mapping of cancer cell states at unprecedented scale and resolution. Single-cell RNA sequencing (scRNA-seq) studies indicate that the constellations of cancer cell states within solid tumors are remarkably reproducible across patients^5^. For instance, in pancreatic ductal adenocarcinoma (PDAC) cancer cells adopt classical, basal, and squamous/mesenchymal identities, which recur across patients^6–9^. Recent meta-analyses of scRNA-seq profiling studies have described a core set of cancer cell differentiation states (“archetype states”) with features that are conserved across different solid tumor types^5,10,11^. Genetically engineered mouse models (GEMMs) of cancer have revealed the pattern in which cancer cell states emerge during tumor evolution is highly stereotypic and ordered^12,13^. These findings imply a recurrence of distinct cancer cell states within tumor tissues that reflects organization of differentiated cells in healthy epithelial tissues, yet this possibility has been little studied. Furthermore, it is not known whether cell state recurrence reflects cooperative relationships between functionally specialized cancer cell states.

The WNT and Notch signaling pathways are among the most prominent molecular mechanisms of tissue pattern generation during development. They also play a key role in the maintenance of epithelial ASCs and in instructing the differentiation of ASC progeny. Secreted WNT ligands act at a short range, typically between neighboring cells. Cells that secrete WNT ligands express porcupine (encoded by the *Porcn* gene in mice), an O-acyltransferase that exclusively palmitoylates WNT ligands – a post-translational modification necessary for secretion and binding to the Frizzled/Lrp receptor complex^14^. Activation of Frizzled/Lrp in WNT-responding cells leads to the stabilization and nuclear translocation of β-catenin. In the nucleus, β-catenin associates with TCF/LEF to activate WNT-responsive genes such as *Lgr5* and *Axin2* together with lineage-specific gene programs controlling differentiation and proliferation^14^.

Notch ligands of the delta-like ligand (DLL) and Jagged families are tethered to the membrane of cells sending Notch signals, requiring direct cell-cell contact for activation of Notch receptors. Furthermore, activation of Notch typically represses Notch ligand expression in signal-receiving cells (lateral inhibition). These properties make the Notch pathway ideally suited for generating patterns in tissues, in which the density of one cell type directly influences the density of other cell types^15^. Downstream signaling drives expression of Notch-responsive genes such as *Hes1* and *Hey1*^15^. In adult epithelia, Notch signaling maintains stem cells and instructs the fate of differentiated cells^15^. WNT inhibition shows promise in PDAC models harboring *RSPO* or *RNF43* mutations^16,17^ as well as in augmenting anti-tumor immunity^18^. However, the combined role of WNT and Notch signaling in PDAC cell fate decisions and tissue organization has not been investigated.

*In vitro* culture methods do not fully recapitulate cancer cell states of human PDAC tumors^9,19^. Thus, studying cell state drivers requires the use of models that produce the full spectrum of cancer and stromal cell diversity in human PDAC. ∼93% of PDACs harbor mutations in the *KRAS* proto-oncogene, whereas the tumor suppressor *TP53* or components of the p53 pathway are mutated in 78% of cases^20,21^. In widely used autochthonous GEMMs of PDAC, expression of ***<u>C</u>****re* recombinase is directed to pancreatic epithelium, inducing somatic activation of oncogenic ***<u>K</u>****ras^G12D^* and mutation of *Tr**<u>p</u>**53* (hereafter “***KPC***” mice), which leads to PDAC development^22–24^. We recently demonstrated that orthotopic allografts of *KPC* cell lines recapitulate the heterogeneity of autochthonous mouse and human PDAC tumors^9^, implying the existence of organizing principles that define PDAC growth *in vivo*.

Emerging evidence indicates distinct cancer cell states manifest in surprisingly reproducible densities and patterns across tumors, implying that malignant cell state heterogeneity in tumors may be significantly more ordered than previously thought^5–9,11,25^. Taking this one step further, cell state patterns in tumors may serve an evolutionary purpose. Here, we employ *KPC* models and patient tissue samples to investigate malignant cell state organization in PDAC, a devastating and intractable disease.

## Results

### PDAC organizes into interdependent WNT-secreting and WNT-responding cancer cell subpopulations

We examined gene expression patterns in distinct cancer cell states in autochthonous genetically engineered *Kras^LSL-G12D/+^; Trp53^flox/flox^; Pdx1-Cre; Rosa26^tdTomato/tdTomato^* (*KPCT*) mouse PDAC tumors using scRNA-seq (**Figure 1a**)^9^. *Porcupine* expression was enriched in classical cancer cells, whereas a subset of basal cells expressed the WNT/β-catenin target *Lgr5* (**Figure 1a**). Porcupine^+^ cancer cells and cells expressing an Lgr5-GFP reporter resided in close proximity in PDAC tumors (**Figure 1b**), consistent with paracrine WNT signaling interactions.

**Figure 1.**
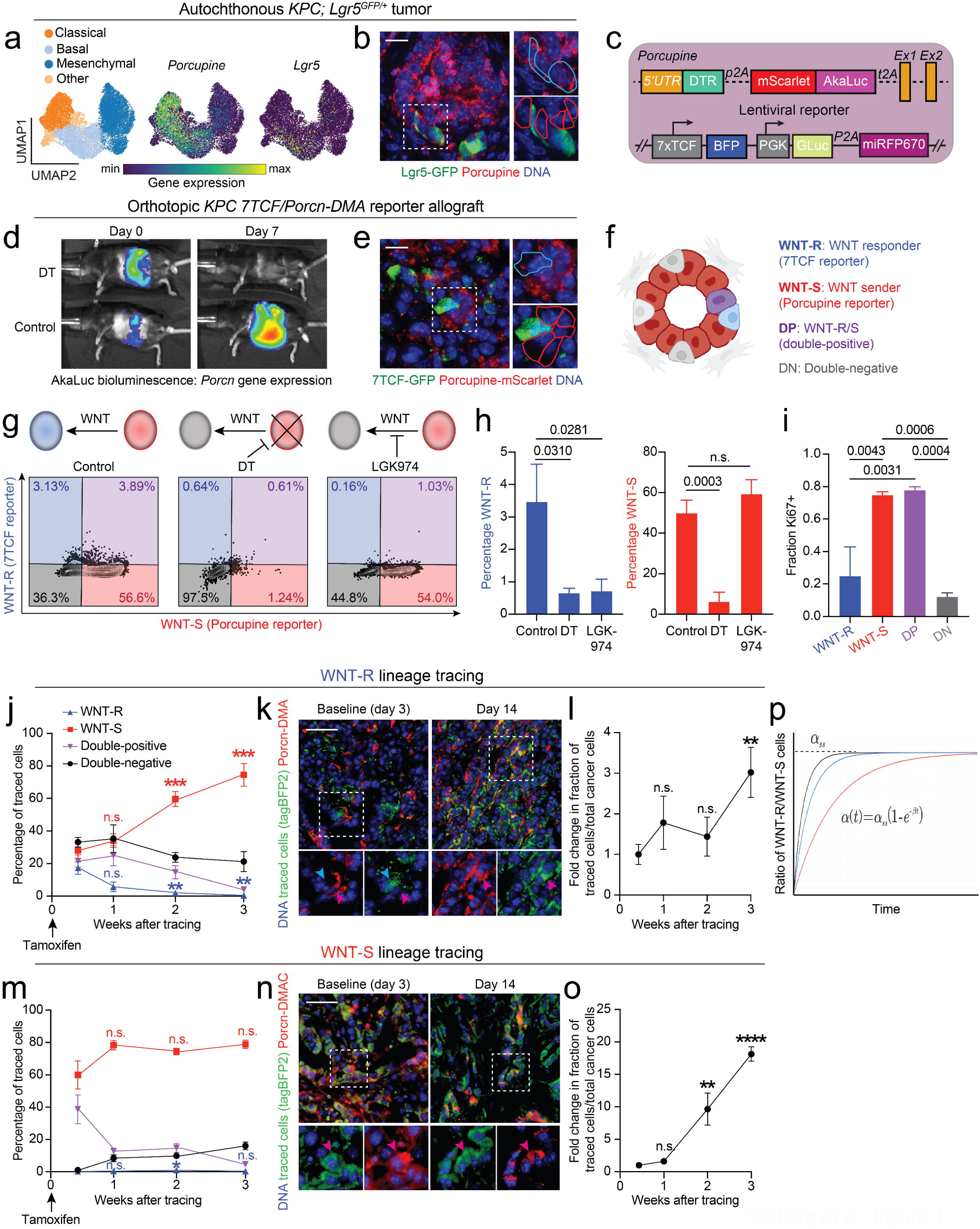
PDAC organizes into interdependent WNT-secreting and WNT-responding cancer cell subpopulations. (**a**) Uniform manifold approximation and projection (UMAP) embedding of scRNA-seq profiles of 14,392 cells from 15 independent *Kras^LSL-G12D/+^; Trp53^flox/flox^; Pdx1-Cre; Rosa26^tdTomato/tdTomato^* (*KPCT*) mouse PDAC tumors, classified as classical (orange), basal (light blue), or mesenchymal (dark blue) cells according to Pitter et al.^9^ (left). *Porcn* (middle) and *Lgr5* (right) expression projected into UMAP. (**b**) Immunofluorescence detection of GFP (green) and porcupine (red) in an autochthonous *KPCT; Lgr5^GFP-CreER/+^* PDAC tumor at 8 weeks. Colored outlines indicate WNT-S (red) and WNT-R (blue) cells. Scale bar: 50 µm. (**c**) Genetic reporters for identifying WNT-S and WNT-R populations engineered into *KPC* cell lines. The *Porcn-DMA* reporter labels porcupine^+^ WNT-S cells in red with mScarlet fluorescence, AkaLuc bioluminescence, and diphtheria toxin receptor (DTR) expression (“*DMA*” cassette, top). The DMA cassette is linked to the endogenous porcupine gene via a P2A peptide. The lentiviral 7TCF reporter labels WNT-R cells with tagBFP2 blue fluorescence (BFP). All tumor cells express *Gaussia* luciferase and the far-red fluorescent protein iRFP670, driven by a ubiquitously active PGK promoter (bottom). (**d**) *In vivo* AkaLuc luciferase bioluminescence imaging of mice with orthotopically transplanted *Porcn-DMA* reporter PDAC tumors at baseline (3 weeks post-transplantation) and after 7 days of diphtheria toxin (DT) administration. Note loss of AkaLuc signal in response to DT, indicating loss of the porcupine^+^ WNT-S cells. (**e**) GFP immunofluorescence (green) and detection of endogenous mScarlet (red) in an orthotopically transplanted *7TCF-eGFP/Porcn-DMA* reporter PDAC tumor. Colored outlines indicate WNT-S (red) and WNT-R (blue) cells. Scale bar: 50 µm. (**f**) Graphical representation of the spatial organization of WNT-S and WNT-R cells. WNT-R cells are a small percentage of tumor cells and reside adjacent to WNT-S cells. (**g**) Representative flow cytometry plots of the WNT-S and WNT-R populations in control, DT-treated, and LGK974-treated orthotopically transplanted *7TCF/Porcn-DMA* reporter PDAC tumors. (**h**) Quantification of WNT-R (blue) and WNT-S (red) subpopulations in experiment described in (**g**); *n* = 7 mice/group. (**i**) Quantification of Ki67+ fraction in WNT-R, WNT-S, WNT-R/S (double-positive, DP), and double-negative (DN) subpopulations in *7TCF/Porcn-DMA* reporter PDAC tumors; *n* = 3 mice. (**j-l**) Lineage-tracing of WNT-R cells in orthotopic *7TCF-CreER/Porcn-DMA* reporter PDAC allografts. (**j**) Quantification of WNT-R, WNT-S, WNT-R/S, and DN populations within the labeled pool of cells following lineage-tracing of the WNT-R cells; *n* = 4-10 mice/time point. (**k**) Representative immunofluorescence images showing tagBFP2-FLAG^+^ traced cells (green) and endogenous mScarlet (red, *Porcn-DMA*) at baseline (3 days) and 14 days after a single pulse of tamoxifen. Blue arrowheads indicate a traced WNT-R, porcupine^−^ cell; red arrowheads indicate WNT-S, porcupine^+^ cells. Note that tagBFP2-FLAG+ traced cells (green) and endogenous mScarlet (red, *Porcn-DMA*) are mutually exclusive at baseline (3 days), whereas they co-localize at 14 days, indicating transdifferentiation of the WNT-R cells to the WNT-S state. (**l**) Fold change in proportion of tagBFP2-FLAG^+^ lineage-traced cells relative to iRFP670^+^ bulk tumor by flow cytometry at distinct time points following labeling of the WNT-R cells. (**m-o**) Lineage-tracing of WNT-S cells in orthotopic *7TCF/Porcn-DMAC* reporter PDAC allografts. (**m**) Quantification of WNT-R, WNT-S, WNT-R/S, and DN populations within the labeled pool of cells following lineage-tracing of the WNT-R cells; *n* = 4-10 mice/time point. (**n**) Representative immunofluorescence images showing tagBFP2-FLAG^+^ traced cells (green) and endogenous mScarlet (red, *Porcn-DMA*) at baseline (3 days) and 14 days after a single pulse of tamoxifen. Red arrowheads indicate WNT-S, porcupine^+^ cells. Note that the tagBFP2-FLAG+ traced cells (green) and endogenous mScarlet (red, *Porcn-DMA*) co-localize at the 3-day and 14-day time points, indicating the WNT-S state is stable. (**o**) Fold change in proportion of tagBFP2-FLAG^+^ lineage-traced cells relative to iRFP670^+^ bulk tumor by flow cytometry at distinct time points following labeling of the WNT-S cells. (**p**) Mathematical modeling demonstrating the ratio of WNT-R and WNT-S cells over time. A pure population of WNT-S cells will establish a consistent equilibrium of WNT-R and WNT-S cells over time. Different slopes represent different values of β. Two-way ANOVA was used in (**h**) and (**i**) to test for statistical significance. Unpaired *t* tests were used to test significance compared to baseline at each time point in (**j**), (**l**), (**m**), and (**o**). Error bars indicate SEM.

To functionally interrogate these porcupine^+^ WNT-secreting (WNT-S) and Lgr5+ WNT-responding (WNT-R) cell states and their interactions, we constructed a multi-component reporter system. We inserted a cDNA cassette linking **<u>d</u>**iphtheria toxin receptor (DTR), **<u>m</u>**Scarlet fluorescent protein, and **<u>A</u>**kaLuc luciferase (***DMA***) into the 5’ untranslated region (UTR) of *Porcn* exon 1 in an established *KPC* cell line (KPC4662)^22,26^ (**Figure 1c; Figure S1a**). We fused the DMA cassette to the endogenous *Porcn* gene using 2A peptide linkage (**Figure 1c**). The targeted insertion of the DMA reporter into the *Porcn* locus did not perturb *Porcn* gene expression (**Figure S1b**) or capacity to secrete active WNT ligands (**Figure S1c**) in the reporter cells. Changes in *Porcn* gene expression correlated with mScarlet fluorescence and AkaLuc bioluminescence in reporter cells subjected to either shRNAs targeting the *Porcn* 3’UTR or *Porcn* overexpression using CRISPR-guided gene activation (CRISPRa) (**Figure S1d, e**). We observed robust AkaLuc bioluminescence upon orthotopic transplantation of the *Porcn-DMA* reporter cells, indicative of *Porcn* gene expression. AkaLuc bioluminescence was extinguished by administration of diphtheria toxin (DT), indicating elimination of the WNT-S cells (**Figure 1d; Figure S1h**). Taken together, these results indicate that *Porcn-DMA* reports for *Porcn* expression at high fidelity without interfering with expression of the endogenous gene and enables ablation of the *Porcn^+^* WNT-S cancer cell state.

To mark the WNT-R cancer cell state, we transduced the *Porcn-DMA* reporter cells with a lentiviral vector comprising a WNT/β-catenin-responsive *7TCF* promoter^27,28^ driving tagBFP2 blue or eGFP green fluorescent protein (**Figure 1c** and not shown, respectively), labeling cells with WNT/β-catenin signaling activity in a second color. The vector also contains a *Gaussia principis* luciferase (G-Luc)-P2A-iRFP670 far-red fluorescent protein cassette placed under the control of a constitutively active phosphoglycerate kinase-1 (*PGK1*) promoter to mark all tumor cells in a third fluorescent color. *Gaussia* luciferase is secreted into the bloodstream, enabling longitudinal tracking of tumor burden via serial sampling of the blood in tumor-bearing mice at high fidelity (**Figure S1f**). The *7TCF/Porcn-DMA* (WNT-R/WNT-S) reporter cells showed no activity of the *7TCF* reporter and moderate, uniform activity of the *Porcn-DMA* reporter in 2D cell culture (**Figure S1d, g**). However, upon orthotopic transplantation the *7TCF/Porcn-DMA* reporter cells organized in a pattern that closely resembled the porcupine and Lgr5-GFP immunostaining pattern observed in the autochthonous *KPC* tumors (**Figure 1b, e, f**). Flow cytometry revealed 3.5% (±3.0%) of the cancer cells reside in the WNT-R state and 50.1% (±16.5%) reside in the WNT-S state (**Figure 1g, h**). WNT-R/S double-positive (DP) cells comprise 2.3% (±2.3%) of cancer cells, whereas double-negative (DN) cells comprise 48.8% (±16.8%) of cancer cells (**Figure 1g, h**). To investigate the fidelity of our reporters *in vivo*, we performed mRNA sequencing (RNA-seq) of WNT-R, WNT-S, WNT-R/S double-positive, and double-negative cancer cell subsets isolated from orthotopic tumor allografts. As expected, the WNT-R cells showed high enrichment of the WNT/β-catenin targets *Lgr5* and *Axin2*, whereas the WNT-S cells were enriched for *Porcn* expression and the WNT-R/S double-positive cells expressed intermediate levels of these genes (**Figure S1i, Table S1**).

To examine the function of the WNT-S cells in PDAC, we ablated them by administering DT to mice with established orthotopic allografts, which led to a 96.6% reduction in the WNT-S population (**Figure 1d, g; Figure S1h**). The fraction of single-positive WNT-R cells was reduced by 65.3% following DT, despite these cells lacking DTR expression (**Figure 1g, h**). Furthermore, targeting ligand-dependent WNT signaling with LGK974^17^, a small molecule inhibitor of porcupine enzymatic activity, produced a 91.8% reduction in the WNT-R population (**Figure 1g, h**). These results indicate WNT ligands produced by the emergence and maintenance of the WNT-R cancer cell subset depends on WNT signals produced by the WNT-S subset.

Notably, 61.2% (±19.8) of the WNT-R cells also showed activity of the WNT-S reporter (**Figure 1g**), implying a transition between these two cell states. Furthermore, the WNT-S and WNT-R/S double-positive cells were highly proliferative in contrast to the WNT-R and double-negative cells (**Figure 1i**), suggesting a significant difference in growth capacity between the cell states. To investigate the fate and clonal expansion potential of the WNT-R and WNT-S cancer cell states *in vivo*, we performed lineage-tracing – the timed introduction of a heritable genetic mark (in our case, a Cre-inducible tagBFP2 fluorescent protein) into a population of interest. To trace the WNT-R cells, we introduced tamoxifen-activatable Cre recombinase (CreER) placed under the control of the *7TCF* promoter (*7TCF-CreER*) and Cre-sensitive “flexed” tagBFP2 into the *7TCF-eGFP*/*Porcn-DMA* cells. In this system, cells in the WNT-R state are marked by GFP, WNT-S cells express mScarlet, and all cancer cells express iRFP670. Upon administration of a single pulse of tamoxifen, cancer cells in the WNT-R state switch on constitutively active tagBFP2 fluorescence (**Figure S2a**). We traced the WNT-R cells in established orthotopic *7TCF-eGFP-CreER*/*Porcn-DMA* reporter allografts and observed a gradual trans-differentiation of the labeled cells towards the WNT-S state (**Figure 1j, k**). The traced cells rapidly exited the WNT-R state (**Figure 1j; Figure S2b**), which was accompanied by clonal expansion of the WNT-R state descendants (**Figure 1l**). These results indicate that (i) the WNT-R state is transient and fated to acquire the WNT-S state and (ii) the WNT-R to WNT-S transition is coupled to an increase in proliferation (**Figure 1i-l**).

To trace the WNT-S cells, we knocked a ***<u>D</u>****TR-**<u>m</u>**Scarlet-**<u>A</u>**kaLuc-**<u>C</u>**reER* (***DMAC***) cassette into the *Porcn* 3’UTR (**Figure S2c, d**). Like the *Porcn-DMA* reporter, the *Porcn-DMAC* system labeled *Porcn*^+^ cells with high fidelity without disrupting gene function (not shown). As expected, both the WNT-S cells and the double-positive WNT-R/S cells were labelled by tagBFP2 in the orthotopic allografts 3 days after the tamoxifen pulse (baseline). The traced WNT-R/S cells rapidly lost activity of the 7TCF-eGFP reporter and acquired the WNT-S single-positive state, which remained stable throughout the 3-week tracing period (**Figure 1m, n; Figure S2e**). Even though the relative proportion of the WNT-S cells remained stable (**Figure S2f)**, we noted that a small fraction of the traced cells acquired a double-negative fate at the longest tracing time points (**Figure 1m, n; Figure S2e**). The traced WNT-S cells exhibited robust clonal expansion (**Figure 1o**), consistent with the high proliferation rate of this cell state (**Figure 1i**).

To account for the dynamics of cancer cell subpopulations and explore potential importance of WNT-R cells in tumor expansion, we developed two simple mathematical models (**Fig. 1p; Supplementary Modeling File 1**). Both models assumed rapid proliferation dynamics of WNT-S cells (with double-negative and double-positive cells grouped with WNT-S cells for simplicity) vs. WNT-R cells, emergence of WNT-R cells from WNT-S cells (or more accurately double negative cells emerging from the WNT-S cells and lumped with them in this model) and rapid conversion of WNT-R cells into WNT-S cells (by way of the intermediate double positive state). The models differed in their assumptions about dependence of WNT-S cell growth on WNT-R cells. In the first model, we assumed that WNT signaling (and thus WNT-R cells) is important for formation of malignant structural units of cell organization and continued WNT-S expansion through increasing number of these units. In the second model, we assumed that WNT-S cells grow independently of WNT-R cells. The models agreed in some predictions and differed in others, allowing us to test the predictions and their mechanistic consequences, and to differentiate between the mechanisms described by the models.

The key consensus prediction by both models was that the fractions of different cell states would stabilize over time to constant levels not dependent on the initial fractions. This prediction was indeed confirmed both by the analysis of the dynamics of cell states. This result provided further evidence for the core of the mechanism postulating interdependence between the WNT-S and WNT-R states, whereby the WNT-S cells induce a transitory WNT-R state in a subpopulation of cells via WNT signals. The WNT-R cells first give rise to the WNT-R/S cells, which subsequently transition into the highly stable and proliferative WNT-S state.

### The WNT-S state and porcupine drive PDAC growth and metastasis

The interdependence and clonal expansion dynamics revealed by lineage tracing of WNT-R and WNT-S cells implies that interactions between these two states drive PDAC growth. At the same time, given the transient nature, relatively low growth rate and low abundance of WNT-R cells, their role in tumor expansion is not clear. The mathematical models varied in their prediction of the growth rates of WNT-S cells constituting the majority of tumor cells. The first model predicted that the growth rate would strongly depend on the ratio of WNT-R/WNT-S cells in the population, whereas according to the second model, this rate is essentially independent of the WNT-R cell number. To experimentally test this, we isolated primary WNT-R, WNT-S, and double-negative (DN) PDAC cells from established orthotopic *7TCF/Porcn-DMA* reporter cell allografts and initiated homotypic and mixed 3D tumor sphere cultures (**Figure 2a**). Mixed cultures of WNT-R and WNT-S cells exhibited considerably higher growth potential compared to all homotypic cultures or the DN + WNT-S mixture (**Figure 2a, b**). These results suggest the WNT-R and WNT-S states cooperate to promote PDAC growth, providing support for the first model and the associated mechanism.

**Figure 2.**
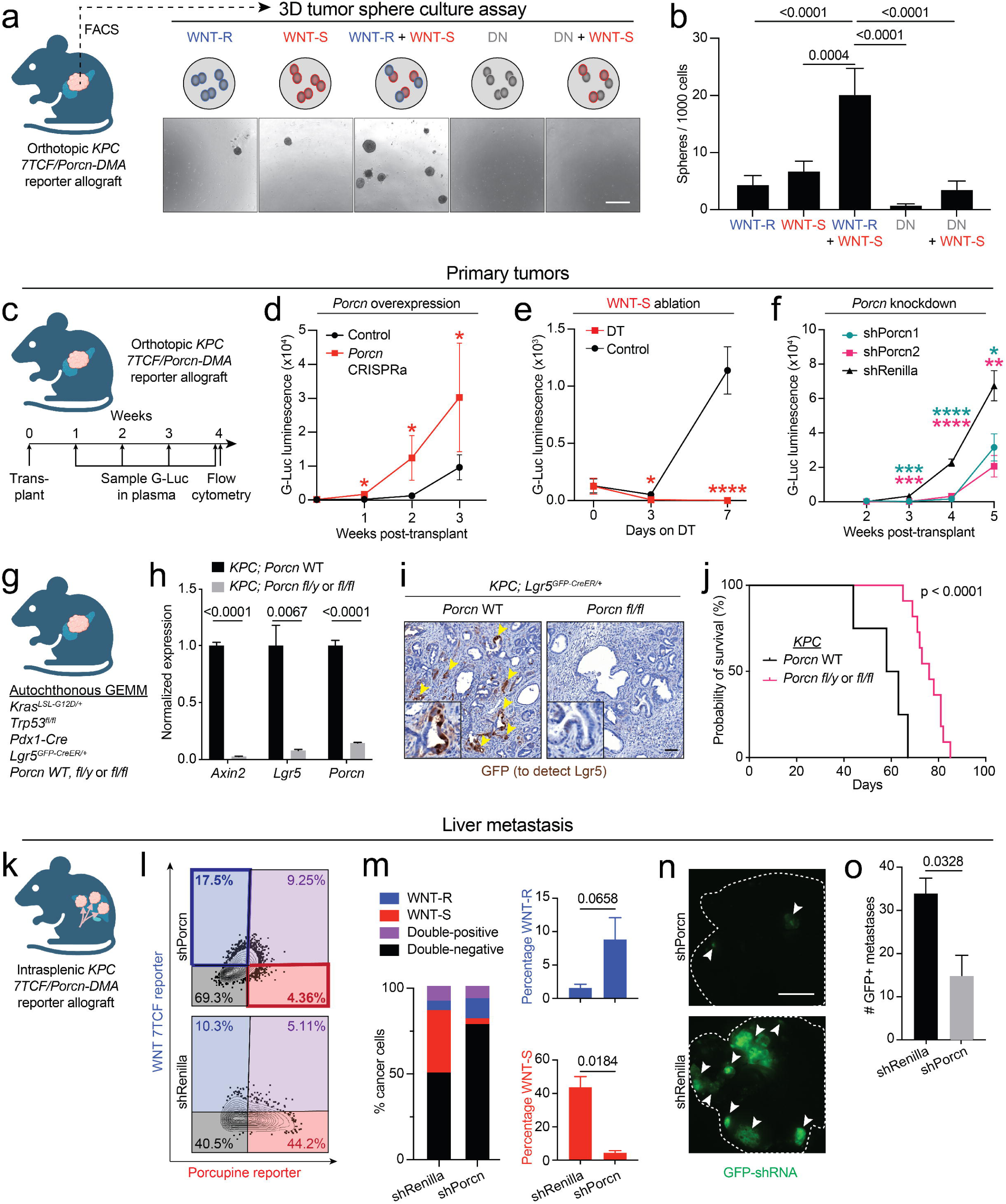
The WNT-S state and porcupine drive PDAC growth and metastasis. (**a**) Experimental design for culturing FACS-isolated WNT-S, WNT-R, and DN cells in 3D tumor sphere culture (top). Representative brightfield images of 3D tumor spheres (bottom). Scale bar: 100 µm. (**b**) Quantification of spheres formed in experiment described in (**a**); *n* = 4 mice, tumor spheres were plated in technical quadruplicates. (**c**) Experimental design for tracking growth of orthotopic *7TCF/Porcn-DMA* reporter PDAC allografts by serial *Gaussia* luciferase (G-Luc) measurements. (**d-f**) Tumor burden assessed by circulating G-Luc in response to *Porcn* overexpression by CRISPRa (*n* = 5-12 mice/group) (**d**); WNT-S state ablation by diphtheria toxin (DT) administration (*n* = 8-9 mice/group) (**e**); or *Porcn* knockdown by shRNA (*n* = 6-7 mice/group) (**f**). (**g**) Autochthonous *KPC*; *Lgr5*^GFP-CreER/+^; *Porcn^flox/flox^* mouse model to interrogate *Porcn* function in *KPC* PDAC tumors. (**h**) qPCR for markers for WNT-R cells (*Axin2*, *Lgr5*) and WNT-S cells (*Porcn*) in wild-type or *Porcn* knockout PDAC tumors; *n* = 3 mice/group. (**i**) Immunohistochemical staining for GFP (brown) to detect Lgr5 in autochthonous *Porcn* wild-type (left) or *Porcn* knockout (right) PDAC tumors. Yellow arrowheads indicate Lgr5^+^ cells. (**j**) Survival curve of mice with autochthonous *KPC* PDAC *Porcn* wild-type or *Porcn* knockout tumors. (**k**) Introduction of liver metastases by intrasplenic injection of *7TCF/Porcn-DMA* reporter PDAC cells. (**l**) Representative flow cytometry plots of *7TCF/Porcn-DMA* reporter PDAC liver metastases in response to *Porcn* or *Renilla* (control) shRNA. (**m**) Left: Stacked bar graph showing percentage of WNT-S, WNT-R, WNT-R/S, and DN cells in *7TCF/Porcn-DMA* reporter PDAC liver metastases from (**l**). Right: Quantification of WNT-R subpopulation (top right) and WNT-S subpopulation (bottom right) fractions in the iRFP670^+^ bulk pool of cancer cells; *n* = 4-10 mice/group. (**n**) Representative whole-liver images of liver metastasis tumors from (**l**) detecting GFP fluorescence in tumors with active shRNA expression. White arrowheads indicate metastatic nodules with GFP expression. White dotted line indicates liver outline traced from brightfield image. Scale bar: 10 mm. (**o**) Quantification of GFP+ metastatic nodules from (**n**). Two-way ANOVA was used in (**b**) to test for statistical significance. Unpaired *t* tests were used to test significance at each time point in (**d**), (**e**), and (**f**). Unpaired *t* tests were used to test significance in (**h**), (**m**), and (**o**). Error bars indicate SEM.

We next evaluated the role of ligand-dependent WNT signaling in the propagation of primary PDAC tumors *in vivo* using orthotopic *7TCF/Porcn-DMA* reporter cell allografts (**Figure 2c**). CRISPRa-mediated overexpression of *Porcn* accelerated tumor growth (**Figure 2d**), whereas DT ablation of the WNT-S state or knockdown of *Porcn* robustly suppressed growth (**Figure 2e, f; Figure S3a**). The first mathematical model accounted for these findings by assuming an increased rate of conversion of WNT-R to WNT-S cells (a prediction validated below by experimental data). Given the experimental data, the model specifically predicted an increase in the exponential growth rate of 0.37/week, due to *Porcn* CRISPRa-mediated overexpression. To further validate these findings in autochthonous PDAC tumors, we crossed a *Porcn^flox/flox^* allele into the *KPC; Lgr5-GFP* PDAC model (**Figure 2g**). Deletion of *Porcn* in the PDAC cells led to near-complete suppression of the WNT targets *Axin2* and *Lgr5* (**Figure 2h**), suggesting that the cancer cells rather than the stroma are the source of WNT ligands that activate WNT/β-catenin signaling in PDAC tumors. As in the allografts, inactivation of *Porcn* resulted in loss of the Lgr5^+^ PDAC cells in the autochthonous *KPC* tumors (**Figure 2i**), suggesting elimination of the WNT-R cell state. Finally, the mice bearing autochthonous *KPC* tumors deficient in *Porcn* survived 26% longer than the mice with *Porcn* WT tumors (**Figure 2j**). These results elucidate a central role for WNT signals produced by the WNT-S cancer cell state in the growth of primary PDAC tumors.

Given that the majority of PDAC patients eventually develop metastatic disease, we next examined cell state organization in liver metastasis – the most common distant-organ site of PDAC dissemination (**Figure 2k**)^29,30^. *7TCF/Porcn-DMA* reporter cells introduced into the liver via intrasplenic injection formed metastases that exhibited highly similar cell state frequencies as the tumors at the primary site (**Figure 1h** vs. **Figure 2l, m**). Similar results were observed in subcutaneous transplants (**Figure S3b)**. These findings suggest PDAC cell state organization is primarily dictated by interactions between malignant cell subsets rather than the tissue site. *Porcn* knockdown suppressed liver metastasis (**Figure 2n, o**), consistent with a central role for cooperative WNT signaling interactions among cancer cells in PDAC dissemination.

### Selective pressure towards equilibrium between WNT-R and WNT-S cell states in PDAC

We next sought to understand how *Porcn* gain-of-function or loss-of-function impact PDAC growth and heterogeneity. We established orthotopic tumors from *7TCF-tagBFP2/Porcn-DMA* reporter cells harboring GFP-linked shRNA or *Porcn* CRISPRa (**Figure 3a; Figure S4a, b**). In this system, all cancer cells are marked by iRFP670, whereas WNT-R cells are blue, WNT-S cells are red, and cancer cells expressing either CRISPRa or shRNA are green fluorescent (**Figure 1c**). In these experiments the genetic perturbations are constitutively active (see STAR Methods, **Figure S4a, b**). At 3 weeks following transplantation, approximately 20% of total iRFP670+ cancer cells had silenced the GFP-linked CRISPRa, shPorcn, or shRenilla, used as a control (**Figure 3b**; **Figure S4c**). We next examined the impact of the gene perturbations on PDAC cell state. Cancer cells that maintained shPorcn-mediated *Porcn* loss were significantly more likely to acquire the WNT-R state, whereas cells that regained *Porcn* expression (i.e., silenced shPorcn) were more likely to acquire the WNT-S state (**Figure 3c, d; Figure S4d**). Conversely, cancer cells that silenced CRISPRa-driven *Porcn* overexpression were significantly less likely to acquire the WNT-S state and showed a non-significant trend towards acquiring the WNT-R state (**Figure 3e, f; Figure S4e**). Thus, genetic loss or gain of the WNT-S cell state in a subset of the cancer cells forces cells that silence the genetic perturbation to acquire the opposite cell state. This implies the presence of feedback mechanisms that sense cell state proportions and identity in PDAC tumors.

**Figure 3.**
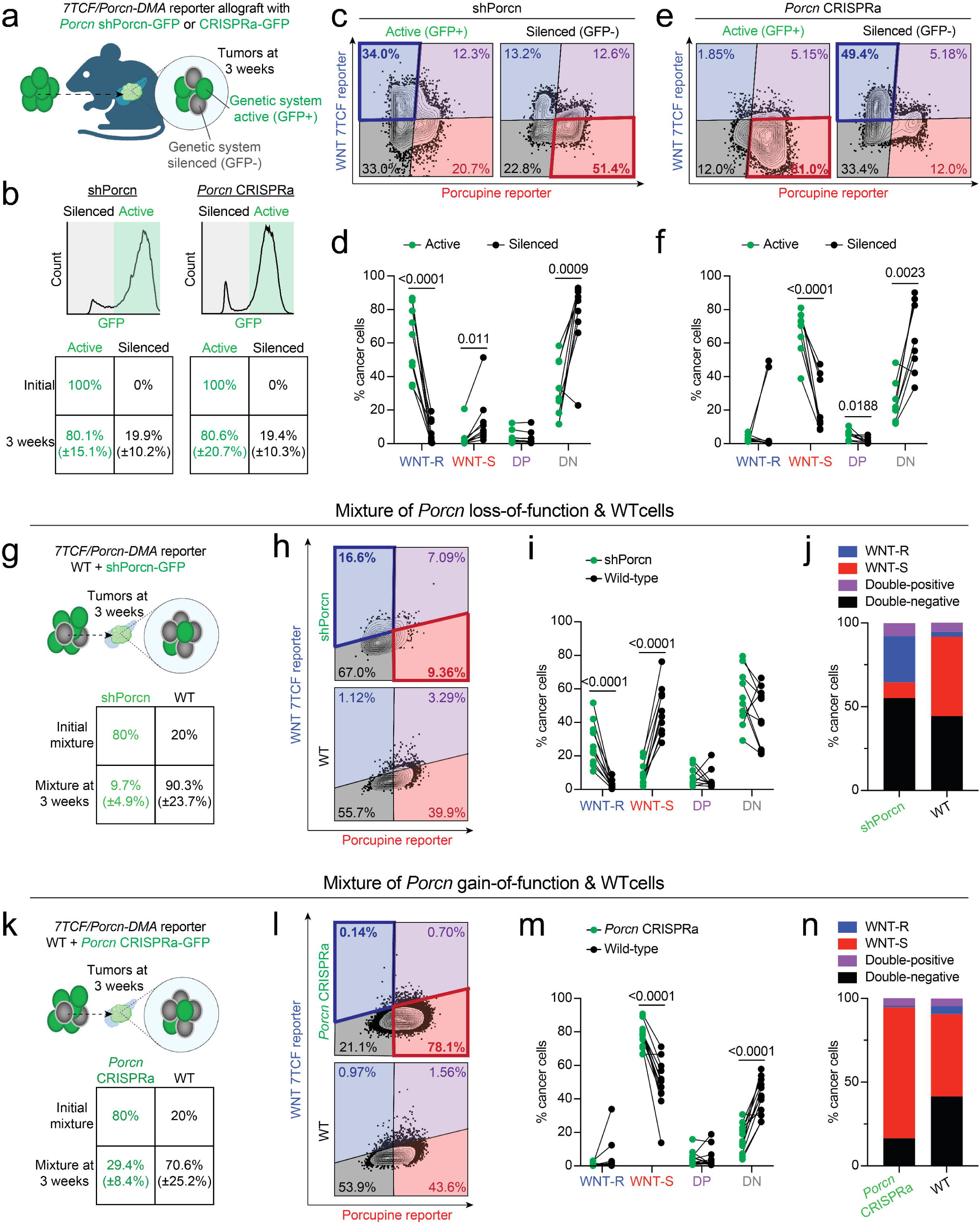
Selective pressure towards equilibrium between WNT-R and WNT-S cell states in PDAC. (**a**) Orthotopic transplantation of *7TCF/Porcn-DMA* reporter PDAC cells constitutively expressing *Porcn* shRNA-GFP or CRISPRa-GFP. Inset: a subset of cells silence either genetic system, as indicated by loss of GFP expression. (**b**) Representative flow cytometry histogram of GFP expression in *Porcn* shRNA-GFP or CRISPRa-GFP tumors (top) and quantification of cells with the genetic system active (GFP+) or silenced (GFP-) initially and at tumor harvest (bottom). The total pool of cancer cells is marked by iRFP670 expression, which was used for gating in flow cytometry (not shown). (**c, e**) Representative flow cytometry plots of orthotopic *7TCF/Porcn-DMA* reporter PDAC allografts harboring shRNA-GFP (**c**) or CRISPRa-GFP (**e**) tumors. Distribution of WNT-S and WNT-R populations in cells with active (left) or silenced (right) genetic constructs. (**d, f**) Quantification of cell states by flow cytometry in shRNA-GFP (**d**) or CRISPRa-GFP (**f**) tumors. Cell populations with active (green dots) or silenced (black dots) genetic constructs from the same tumor are connected by a line; *n* = 5-7 mice/group. (**g**) Orthotopic transplantation of mixed populations of wild-type or *Porcn* shRNA-GFP^+^ *7TCF/Porcn-DMA* reporter PDAC cells (top). 80% of the transplanted cells express *Porcn* shRNA-GFP and 20% of the cells are wild-type at the start of the experiment. Quantification of *Porcn* shRNA-GFP^+^ vs. wild-type cells at 3 weeks post-transplantation is shown in the bottom row of the foursquare plot. (**h**) Representative flow cytometry plots of cell states in *Porcn* shRNA-GFP^+^ and wild-type populations in an orthotopic mixed *7TCF/Porcn-DMA* reporter allograft. Tumor cells are gated to show the distribution of WNT-S and WNT-R populations in cells with active genetic constructs (top) or wild type cells (bottom). (**i**) Quantification of cell states in experiment described in (**g**) and (**h**). Cell populations with genetic constructs (green dots) or wild type cells (black dots) in the same tumor are connected by a line; *n* = 10-12 mice/group. (**j**) Stacked bar graph showing the percentage of WNT-S, WNT-R, WNT-R/S, and DN cells in a *Porcn* shRNA-GFP^+^ and wild-type mixed allograft. (**k**) Orthotopic transplantation of mixed populations of wild-type or *Porcn* CRISPRa-GFP^+^ *7TCF/Porcn-DMA* reporter PDAC cells (top). 80% of the transplanted cells express *Porcn* CRISPRa-GFP and 20% of the cells are wild-type at the start of the experiment. Quantification of *Porcn* CRISPRa-GFP^+^ vs. wild-type cells at 3 weeks post-transplantation is shown in the bottom row of the foursquare plot. (**l**) Representative flow cytometry plots of cell states in *Porcn* CRISPRa-GFP^+^ and wild-type populations in an orthotopic mixed *7TCF/Porcn-DMA* reporter allograft. Tumor cells are gated to show the distribution of WNT-S and WNT-R populations in cells with active genetic constructs (top) or wild type cells (bottom). (**m**) Quantification of cell states in experiment described in (**k**) and (**l**). Cell populations with genetic constructs (green dots) or wild type cells (black dots) in the same tumor are connected by a line; *n* = 10-12 mice/group. (**n**) Stacked bar graph showing the percentage of WNT-S, WNT-R, WNT-R/S, and DN cells in a *Porcn* CRISPRa-GFP^+^ and wild-type mixed allograft. Chi-squared test was used in (**b**), (**g**), and (**k**) to test for statistical significance. Two-way ANOVA was used in (**d**), (**f**), (**i**), and (**m**) to test for statistical significance. Values in parentheses in (**b**), (**g**), and (**k**) indicate SD.

Our experimental results and modeling suggest that PDAC tumors favor an equilibrium of WNT-R and WNT-S cell states for optimal growth. To directly test this possibility, we mixed *7TCF-tagBFP2/Porcn-DMA* reporter cells harboring GFP-linked shRNA targeting *Porcn* or *Renilla* with wild-type (WT) *7TCF-tagBFP2/Porcn-DMA* reporter cells lacking shRNA constructs at an 80:20 ratio (**Figure 3g**). At 3 weeks post-transplantation the WT cells had expanded significantly more than the cells expressing either *Porcn* or *Renilla* shRNA (**Figure 3g**; **Figure S4f, g**). As in the previous experiment, cancer cells expressing shPorcn were significantly more likely to acquire the WNT-R state and maintained low expression of the *Porcn-DMA* reporter (**Figure 3h-j**). No differences in cell state acquisition were observed between shRenilla and WT cells (**Figure S4h**). Further, WT cancer cells expanded significantly more than cells harboring the CRISPRa *Porcn* gain-of-function system (**Figure 3k**). However, the CRISPRa cells exhibited a relative growth advantage when compared to the shPorcn and shRenilla cells (**Figure S4g**), suggesting that maintenance of a subset of cancer cells with high *Porcn* expression is favored over *Porcn* loss-of-function (or neutral control) in the mixed tumors. The *Porcn* CRISPRa cells maintained high *Porcn-DMA* reporter activity and showed a non-significant trend towards lower likelihood of acquiring the WNT-R state when compared to the WT cells (**Figure 3l-n)**. Taken together, these results suggest a selective pressure towards maintenance of an equilibrium between WNT-R and WNT-S cell states in PDAC.

### Notch signaling controls equilibrium of WNT-S and WNT-R states in PDAC

Our findings indicate that the WNT-R state is dependent on WNT signals produced by the WNT-S state (**Figure 1g, h**). However, the frequency of the WNT-S state is 5-to 10-fold higher than the WNT-R state and, although all WNT-R cells neighbor WNT-S cells, not all WNT-S cells neighbor WNT-R cells. This suggested that additional signaling mechanisms may be present in a subset of the WNT-S cells that forms the functional niche for WNT-R cells. Given its function in tissue patterning and lateral inhibition, we explored the Notch pathway as a candidate mechanism. We found a subset of the classical *Porcn*^+^ PDAC cells co-express the Notch ligand *Dll1* in autochthonous *KPC* tumors (**Figure 4a, b**). Furthermore, the Lgr5^+^ cancer cells express high levels of the Notch targets hairy and enhancer of split-1 (*Hes1*) and Notch-responsive ankyrin repeat protein (*Nrarp*) (**Figure 4a, b**). We focused on NRARP since it is also a WNT target and integrates Notch and WNT signaling inputs^31,32^. We found the DLL1^+^ cancer cells localize next to NRARP^+^ cancer cells (**Figure 4c; Figure S5a**), suggesting that DLL1 may contribute to NRARP expression in the WNT-R cells.

**Figure 4.**
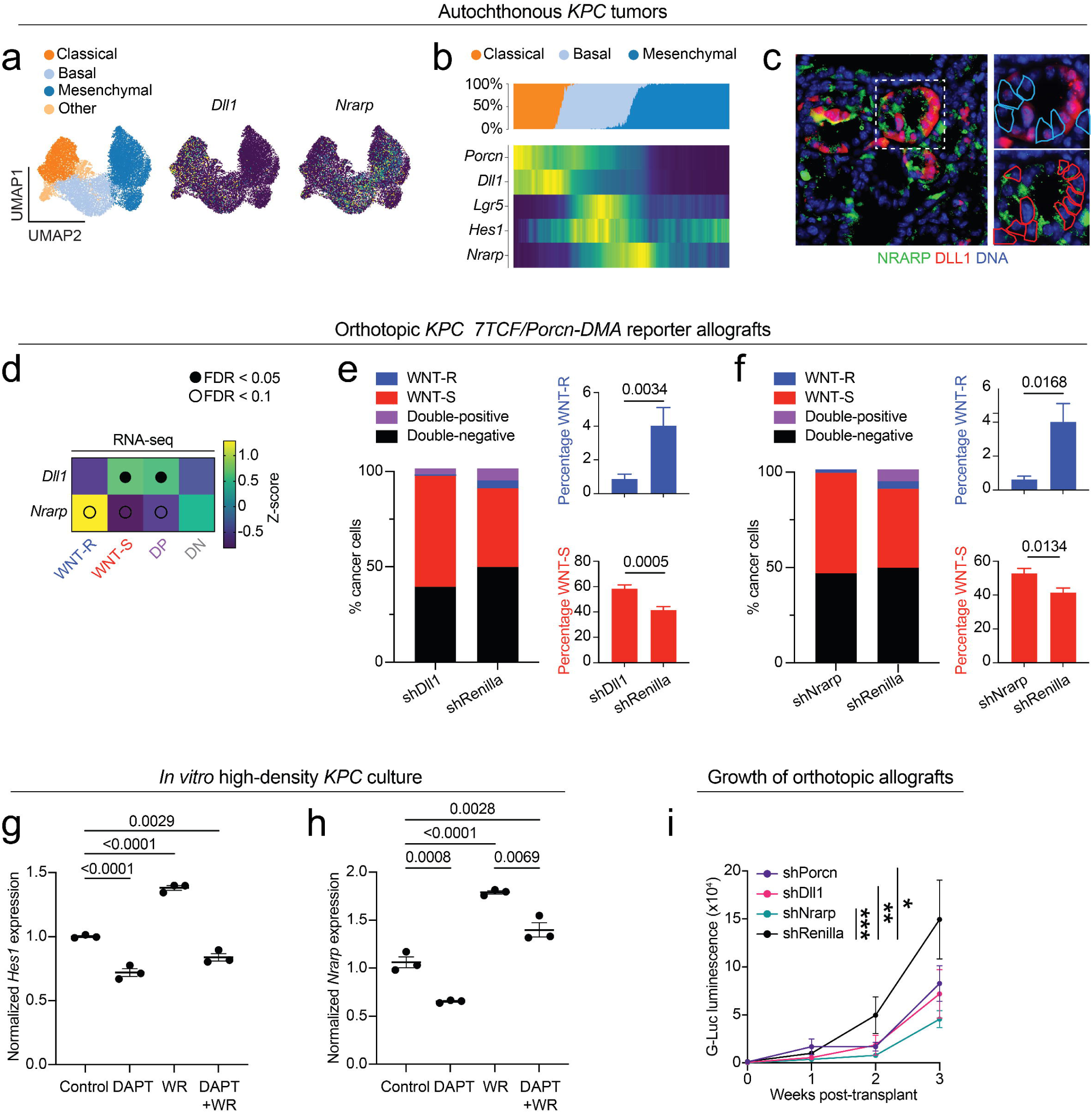
Notch signaling controls equilibrium of WNT-S and WNT-R states in PDAC. (**a**) Uniform manifold approximation and projection (UMAP) embedding of scRNA-seq profiles of 14,392 cells from 15 independent *Kras^LSL-G12D/+^; Trp53^flox/flox^; Pdx1-Cre; Rosa26^tdTomato/tdTomato^* (*KPCT*) mouse PDAC tumors, classified as classical (orange), basal (light blue), or mesenchymal (dark blue) cells according to Pitter et al. (9) (left). *Dll1* (middle) and *Nrarp* (right) expression projected into UMAP. (**b**) *Porcn*, *Dll1*, *Lgr5*, *Hes1*, and *Nrarp* expression *KPCT* cells ordered along the classical-basal-mesenchymal pseudotime axis reported in ref.^9^. Note the expression of *Porcn* and *Dll1* in classical cells and expression of *Lgr5*, *Hes1*, and *Nrarp* in basal cells. Percentage of cells at each pseudotime step classified as classical, basal, or mesenchymal (top). (**c**) NRARP (green) and DLL1 (red) immunofluorescence in an autochthonous *KPCT* PDAC tumor. Colored outlines indicate boundaries of WNT-S (red) and WNT-R (blue) cells. Note that expression is mutually exclusive and NRARP^+^ cells are adjacent to DLL1^+^ cells. Scale bar: 50 µm. (**d**) RNA sequencing (RNA-seq) analysis of *Dll1* and *Nrarp* expression in primary WNT-R, WNT-S, WNT-R/S, and DN cells isolated from orthotopic PDAC *7TCF/Porcn-DMA* reporter allografts. Expression is quantified as a row-normalized z-score; *n* = 4 mice/group. (**e**, **f**) Percentage of WNT-S, WNT-R, WNT-R/S, and DN cells in *7TCF/Porcn-DMA* reporter PDAC allografts in response to shRNA-mediated *Dll1* (**e**) or *Nrarp* knockdown (**f**) vs. Renilla controls; *n* = 7-8 mice/group. (**g**, **h**) *Hes1* (**g**) and *Nrarp* (**h**) expression assessed by quantitative PCR (qPCR) in high density *in vitro* culture of *KPC* cells with WNT3a + R-spondin-1 supplementation (WR), the Notch inhibitor DAPT, or vehicle controls; *n* = 3 technical replicates. (**i**) Tumor burden assessed by circulating G-Luc in response to *Dll1* or *Nrarp* knockdown or shRenilla control in orthotopic *7TCF/Porcn-DMA* reporter PDAC allografts; *n* = 7-8 mice/group. One-way ANOVA with FDR was used in (**d**) to test statistical significance. Open circles indicate FDR < 0.1; closed circles indicate FDR < 0.05. Unpaired *t* tests were used to test significance in (**e**), (**f**). One-way ANOVA was used to test significance in (**g**), (**h**), and (**i**). Error bars indicate SEM.

We next functionally interrogated *Dll1* and *Nrarp* in orthotopic *7TCF-tagBFP2/Porcn-DMA* reporter allografts. Consistent with the autochthonous tumors, we found *Dll1* expression is high in the WNT-S subset, whereas the WNT-R cells express high levels of *Nrarp* (**Figure 4d, Table S1**). Furthermore, knockdown of either *Dll1* or *Nrarp* dramatically suppressed the WNT-R state and increased the frequency of WNT-S cells, as assessed by the *7TCF-tagBFP2* and *Porcn-DMA* reporters (**Figure 4e, f; Figure S5b, c**). These results suggest DLL1-mediated induction of NRARP downstream of Notch is necessary for maintaining the WNT-R state in PDAC tumors. We next examined the cooperation of WNT and Notch signals in *KPC* cells cultured at high density to permit cell contact-dependent Notch ligand-receptor interactions. Stimulation with recombinant <u>W</u>NT3a and <u>R</u>-spondin-1 (WR) promoted *Hes1* and *Nrarp* expression, whereas inhibiting Notch with the γ-secretase inhibitor DAPT suppressed *Hes1* and *Nrarp* both at baseline and in the presence of WR. DAPT suppressed *Hes1* to a similar level both at baseline and in the presence of WR. In contrast, *Nrarp* was only partially suppressed by DAPT in the presence of WR (**Figure 4g, h**). These results demonstrate both WNT and Notch signals are needed for maximal *Nrarp* induction in PDAC cells.

Finally, knockdown of *Nrarp* or *Dll1* suppressed growth of orthotopic PDAC allografts to a similar extent as silencing *Porcn* (**Figure 4i**). Taken together, our findings support a model whereby a subset of WNT-S cancer cells provide WNT ligands and the Notch ligand DLL1, which induce NRARP in adjacent cells, leading to acquisition of the WNT-R state. Each of the components – Porcupine, DLL1, and NRARP – are necessary for inducing the WNT-R state and driving PDAC growth.

### DLL1^+^ WNT-S and NRARP^+^ WNT-R states are conserved in human PDAC

To explore the relevance of our findings in the mouse models to human PDAC, we examined gene expression patterns in The Cancer Genome Atlas (TCGA) and Genotype-Tissue Expression (GTEx) datasets. *PORCN* was significantly induced in PDAC tissues when compared to normal pancreas (**Figure 5a**). Immunostaining of a PDAC tissue microarray comprising 261 patients revealed porcupine localized to malignant cells with a highly epithelial/classical morphology (**Figure 5b**). The majority of patients (56.3%) displayed staining in a subset of PDAC cells (**Figure 5b, c**; **Figure S6**), suggesting competence for WNT secretion in a subpopulation of PDAC cells. To quantify cell state densities at high resolution, we visualized WNT-R and WNT-S cells by *in situ* detection of *LGR5* and *PORCN* mRNA, respectively (**Figure 5d**). We found *LGR5^+^*and *PORCN^+^* cancer cells were present in similar fractions as in the mouse models (3.7% ±0.6% and 37.9% ±13.0%, respectively) (**Figure 5e**). Similarly, a large majority of the *LGR5^+^* cells neighbored the *PORCN^+^* cells (86.3% ±5.1%), whereas *PORCN^+^* cells were not always in contact with the *LGR5^+^* cells (51.7% ±16.7%) (**Figure 5f**). Finally, *DLL1* expression was enriched in a subset of *PORCN^+^* cells, whereas NRARP expression was more frequently observed in *LGR5^+^* cells than in *LGR5^−^* cells (**Figure 5g-j**). These findings suggest that human and mouse PDAC cell state organization follow similar principles (**Figure 5k**).

**Figure 5.**
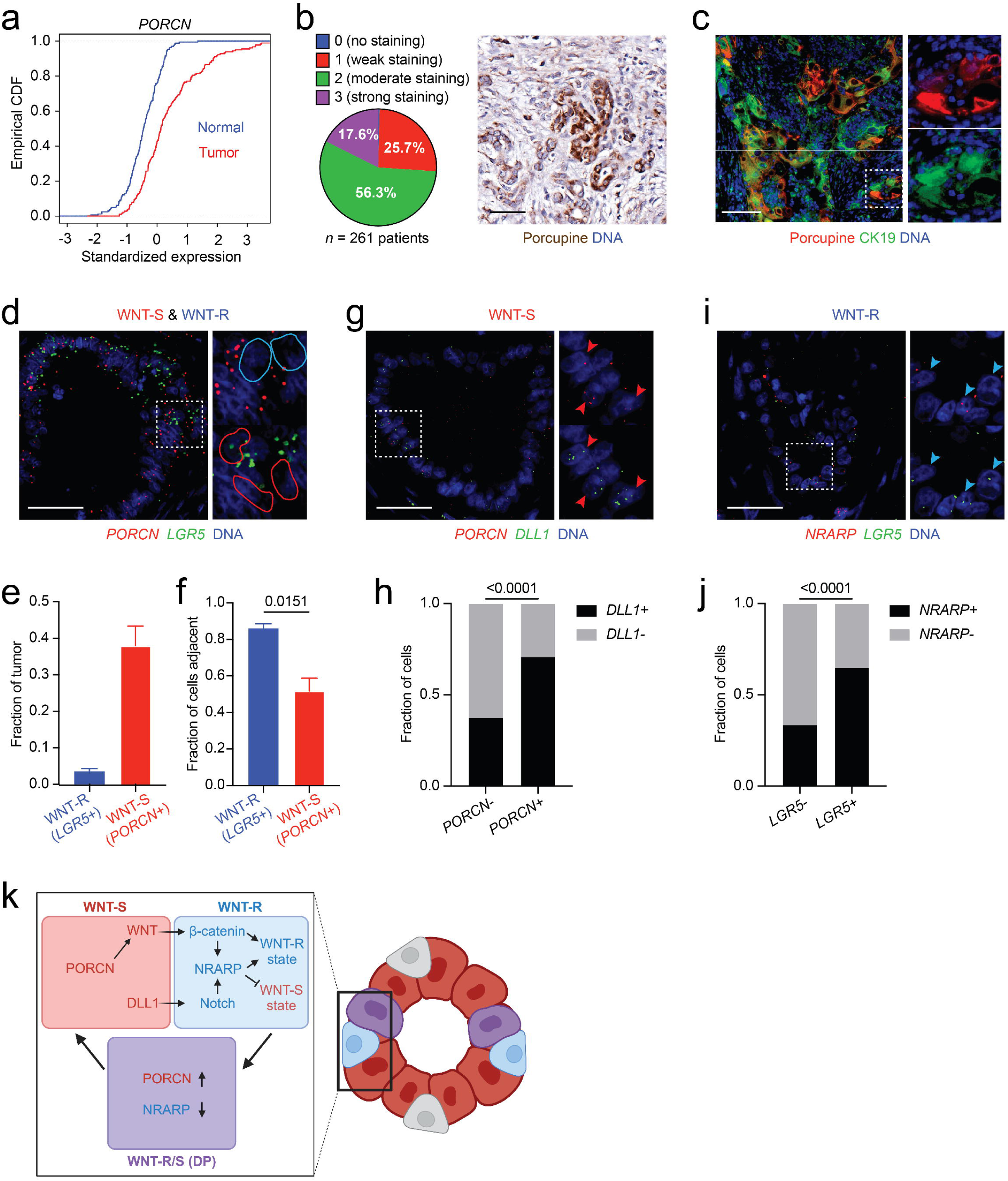
DLL1^+^ WNT-S and NRARP^+^ WNT-R states are conserved in human PDAC. (**a**) *PORCN* expression from bulk RNA-seq derived from the Genotype-Tissue Gene Expression (GTEx) Portal in normal human pancreas (blue) and TCGA human PDAC (red) plotted as an empirical cumulative distribution function (CDF). Rightward shift indicates higher expression in tumors (*p* < 10^−10^ Kolmogorov-Smirnov test); *n* = 150 tumor and 165 normal samples. (**b**) Staining intensity classification (left) and representative image (right) of moderate porcupine staining (brown) in human PDAC tumors, *n* = 261 tumor samples. Scale bar: 50 µm. (**c**) Cytokeratin-19 (CK19, green) and porcupine (red) in a human PDAC tumor. Scale bar: 50 µm. (**d**) RNA *in situ* hybridization of *PORCN* (red) and *LGR5* (green) in human PDAC tissues. Colored outlines indicate boundaries of WNT-S (red) and WNT-R (blue) cells. Scale bar: 50 µm. (**e**) Quantification of proportions of *LGR5^+^* WNT-R cells and *PORCN^+^* WNT-S cells in human PDAC tumors; *n* = 3 patients. (**f**) Quantification of WNT-R cells adjacent to WNT-S cells and WNT-S cells adjacent to WNT-R cells in human PDAC tumors; *n* = 3 patients. (**g**) *PORCN* (red) and *DLL1* (green) RNA *in situ* hybridization in human PDAC. Red arrowheads indicate cells with co-localization of *PORCN* and *DLL1*. (**h**) Quantification of *DLL1* expression in *PORCN^+^*vs *PORCN^−^* cells; *n* = 3 patients. (**i**) *NRARP* (red) and *LGR5* (green) RNA *in situ* hybridization in human PDAC. Blue arrowheads indicate cells with co-localization of *NRARP* and *LGR5*. (**j**) Quantification of *NRARP* expression in *LGR5^+^* vs. *LGR5^−^* cells, *n* = 3 patients. (**k**) Graphical summary of the findings (see text). Kolmogorov-Smirnov test was used to determine statistical significance in (**a**). Unpaired *t* test was used to test significance in (**f**). Chi-squared test was used in (**b**), (**g**), and (**h**) and (**j**) to test for statistical significance. Error bars indicate SEM.

## Discussion

Our findings elucidate the organization of functionally distinct cancer cell differentiation states in tumor tissues. We discover that not only do malignant cell states manifest reproducibly in PDAC tumors, but they also unexpectedly organize in tightly controlled frequencies within the tumor tissue. We find the stereotypic organization of PDAC cell states emerges even in transplants of established cell lines – including single-cell clones derived from such cell lines, as is the case for our *Porcn-DMA* and *Porcn-DMAC* reporter cells – and occurs at both primary and metastatic sites. This indicates malignant cell state identity is controlled by differentiation signals rather than genetic mutations. Our modeling results indicate the lineage and growth dependencies between different cancer cell states produces a consistent equilibrium in the tumor over time. This suggests selective pressure to acquire functionally specialized cell states and establish cooperative relationships during tissue growth is deeply engrained in metazoan cells and not erased by malignant transformation.

Our findings indicate PDAC tissue expands by a stereotypic patterning process, mimicking organogenesis. Our experimental and modeling results are consistent with a model whereby this patterning is orchestrated by DLL1^+^ WNT-S cells, which induce the WNT-R state in a subset of basal PDAC cells. The WNT-R cell state is relatively quiescent and highly transient, rapidly giving rise to WNT-S cells that proliferate and expand to complete the growth cycle – followed by a new cycle (**Figure 5k**). This model suggests that tumor growth *in vivo* is more complicated than previously thought. Rather than all malignant cells exhibiting uniform growth rates, distinct cancer cell states maintain either high or low proliferation rates. Given that the DLL1^+^ WNT-S cells are critical for initiating and directing tissue unit generation, the mechanisms controlling the DLL1^+^ WNT-S state remain an important area of future study. Basal cells give rise to differentiated epithelium during the regeneration of bronchiolar epithelium^33^ and mammary ducts^34^, suggesting our observations may reflect subversion of a similar mechanism in PDAC. Until recently, pancreatic ducts were considered to lack basal cells. However, emerging evidence suggests rare basal cells line large ducts in the human pancreas and may participate in duct regeneration^35^. Further studies are needed to understand the connections between cell states in healthy or regenerating pancreas and the WNT-R and WNT-S states that we uncovered in PDAC tumors.

Oncogenic KRAS is critical for driving PDAC^26,36^, yet our results point to another important layer of growth control that manifests *in vivo*. An equilibrium of cancer cell states, established through differing proliferation rates and lineage interconversion, optimizes growth in cells primed with oncogene activation. Disruption of this equilibrium is deleterious for tumor growth. In the context of PDAC, this growth control is directed by coordinated WNT and Notch signals that instruct malignant cell differentiation, which, in turn, gates KRAS-driven proliferation in PDAC. Specification of pancreatic progenitors from the foregut requires inhibition of WNT signaling, whereas expansion of that tissue is promoted by WNT signaling^37^. In later stages of development, WNT promotes differentiation of pancreatic progenitors into acinar cells^37^. Furthermore, WNT signaling contributes to injury responses in the pancreas: WNT ligands and nuclear β-catenin are required for reacquisition of the acinar fate following acinar-to-ductal metaplasia (ADM) – an essential process in pancreas injury repair^38^. WNT-responsive cancer cell populations harboring directed differentiation potential have been identified in intestinal and lung tumors^39–42^, suggesting similar mechanisms may control cell state organization in other cancers beyond PDAC. Notch ligands and receptors are expressed at various stages of pancreas development, which act via lateral inhibition to specify endocrine differentiation and maintain pancreatic progenitor states^43^. Notably, DLL1 is the primary ligand responsible for Notch activation in this process^43^, similar to our findings in PDAC. Although the adult pancreas shows little Notch activity, signaling is reactivated upon injury: Cerulein-induced pancreatitis induces Notch signaling in ADM, whereas pathway inhibition leads to impaired regeneration and decreases the frequency of mature acinar cells^44,45^. Thus, PDAC cell state organization may mimic pancreas organogenesis and regeneration, which involves WNT and Notch as well as RAS activation downstream of growth factor signals.

The stereotypic patterns of cancer cells in tumors under light microscopy have been used by pathologists for over a century to diagnose and classify tumors. Contemporary methods such as single-cell genomics have greatly expanded our understanding of distinct cancer cell differentiation states in tissues. These high-resolution molecular studies have revealed that cell states within tumors are highly stereotypic and reproducible across individual tumors^5–9,11,25^, reflecting the stereotypic histopathological patterns that define cancers. Initial studies employing spatial genomics or proteomics approaches and innovative computational methods have revealed that distinct cancer cell states reside in defined spatial locations within tumors^46,47^. However, interrogation of the dynamics and molecular mechanisms of cell state organization requires functional studies. Our work highlights the power of combining reporter systems, lineage-tracing, lineage-ablation, and genetic perturbation in the investigation of cell state relationships. Through these approaches we uncovered the WNT-S state as a candidate therapeutic target for cytoablative therapies, whereas DLL1 and NRARP emerged as novel molecular targets in PDAC. Therapeutic strategies targeting the WNT-S state or blocking DLL1 or NRARP function are founded on disruption of cell state organization and heterogeneity within tumors. This contrasts with the traditional oncoprotein-targeted therapies, chemotherapies, or immunotherapies, which have to date shown little efficacy in PDAC. Bold, new therapeutic concepts are urgently needed for PDAC, which portends a 5-year survival of 10% and is on track to become the second-most common cause of cancer-related death.

Expansion by repeating growth cycles where frequency of malignant differentiation states is tightly controlled provides a novel framework for conceptualizing tumor growth. We identify WNT and Notch as central molecular mechanisms underpinning tissue unit assembly in PDAC and show that disrupting the equilibrium of cell states produces a robust anti-tumor response. As such, our work constitutes a departure from the paradigm that tumors are chaotic tissues lacking organization. Instead, we elucidate a highly ordered cell state composition that closely resembles healthy organs. The maintenance of order among cell states is dependent on cooperative interactions and lineage relationships between functionally distinct cell states. Given that most solid tumors show stereotypic patterns of heterogeneity^5,10^, our model of cancer cell state organization and equilibrium as a layer of growth control may represent a more general mechanism by which cancers grow. Targeting the cancer cell state equilibrium may translate into transformative therapeutic strategies.

## STAR Methods

### Animal Studies

All animal studies were approved by the MSKCC Institutional Animal Care and Use Committee (protocol #17-11-008). MSKCC guidelines for the proper and humane use of animals in biomedical research were followed. All genetically engineered mice were maintained on C57/BL6 and Sv129 mixed backgrounds. Previously published autochthonous *Kras^LSL-G12D/+^; Trp53^flox/flox^; Pdx1-Cre; Rosa26^tdTomato/tdTomato^* (*KPCT*); *Lgr5^GFP-IRES-CreER/+^* mice were used in this study^9^. *KPCT*; *Lgr5^GFP-IRES-CreER/+^*; *Porcn ^flox/flox^* mice were generated by crossing *KPCT*; *Lgr5^GFP-IRES-CreER/+^* and *Porcn^flox/flox^* mice^48^. Mouse genotyping was performed at 2 weeks and tumors were harvested at 7-9 weeks when mice demonstrated PDAC symptoms (weight loss, distended abdomen, hunching). Mice were euthanized by CO_2_ asphyxiation followed by intracardiac perfusion with PBS to clear tissues of blood.

### Orthotopic models

Tumor allografts were introduced into wild-type C57/BL6 or immunodeficient NOD.Cg-*Prkdc^scid^; Il2rg^tm1Wjl^*/SzJ (NSG) mice. Murine KPC4662 PDAC cells were obtained from Dr. Robert Vonderheide, generated from a *Kras^LSL-G12D/+^, Trp53^LSL-R172H/+^, Pdx1-Cre* mouse in the C57BL/6J genetic background^22,26^. Cell lines were orthotopically transplanted by injecting 30,000 cells from 2D cell culture into the tail of the pancreas, as before^49^. The presence of tumors was confirmed by trans-abdominal palpation or detection of *Gaussia princeps* luciferase in the blood of the mice. All animals were monitored weekly, and tumors were harvested at a maximum size of 1 cm^3^ or if the animals met other criteria for humane euthanasia.

### Intrasplenic transplantation

Liver metastasis allografts were performed in NOD.Cg-*Prkdc^scid^ Il2rg^tm1Wjl^*/SzJ (NSG) mice. Metastases were generated by intrasplenic injection of 100,000 *KPC 7TCF/Porcn-DMA* cells harboring doxycycline (Dox)-inducible shPorcn-GFP or shRenilla-GFP followed by splenectomy after letting tumor cells circulate for 5 minutes. Mice were maintained on doxycycline hyclate chow (doxycycline 625 mg/kg, Envigo) starting one day before transplantation. Tumor-bearing livers were harvested at 5-6 weeks following transplantation.

### Lineage-tracing

Orthotopic *KPC* cell line allografts harboring WNT-S or WNT-R lineage tracing vectors were established in NSG mice and monitored as above. Tumor-bearing mice were treated with one dose of tamoxifen (200 mg/kg via oral gavage) 1- or 2-weeks post-transplantation. Pancreatic tumors were harvested 3, 7, 14, or 21 days after tamoxifen administration. The baseline measurement at 3 days was chosen to account for conversion of tamoxifen to the active metabolite 4-hydroxytamoxifen (4-OHT), recombination, and expression of the FLEx-tagBFP in lineage traced cells, and elimination of residual 4-OHT. Tamoxifen was dissolved in corn oil at 20 mg/mL and dissolved at 55C for 1 hour, as before^41^.

### Mixed population orthotopic transplantation

Mixed transplants of *7TCF/Porcn-DMA* reporter cells and wild-type PDAC cells harboring *Porcn* shRNA-GFP or CRISPRa-GFP were performed by *in vitro* pooling of 24,000 (80% of 30,000) *Porcn* shRNA-GFP or CRISPRa-GFP cells with 6,000 (20% of 30,000) wild-type cells, followed by orthotopic injection as described above. In the wild type cells, the iRFP670 cassette marking all cancer cells was replaced with Thy1.1 to distinguish the wild-type cells from the cells with the gene perturbation by flow cytometry.

### AkaLuc bioluminescence imaging in vivo

NSG mice bearing orthotopic transplants of *7TCF/Porcn-DMA* reporter cells were dosed with 100 µl of 30 mM of AkaLumine-HCl substrate resuspended in PBS (#808350, Millipore Sigma, TokeOni) and imaged on an IVIS Lumina II (Perkin Elmer)^50^. Signal was recorded for 15 1-minute exposures to capture peak luminescence.

### Plasma sampling and Gaussia princeps luciferase measurements

Whole venous blood was harvested by puncturing the submandibular vein, followed by collection of 100 µl blood into collection vials (#02-675-185, Fisher Scientific)^51^. Plasma was separated by centrifugation at >8000 g for 10 minutes at 4°C and diluted 1:10 in PBS. 200 µM *Gaussia* luciferase substrate coelenterazine-h (#301, NanoLight) was added, and luminescence was immediately measured on a BioTek Cytation 1 (Agilent).

### Gene knockdown in vivo

PDAC cells expressing GFP-linked, doxycycline-inducible shRNAs^52^, along with the *7TCF/Porcn-DMA* reporter constructs were transplanted into NSG mice as described above. Short-term knockdown was induced 3 weeks after transplantation, when the tumors were detectable by *Gaussia* luciferase, prior to onset of tumor-associated symptoms, by placing the NSG mice on doxycycline hyclate chow (doxycycline 625 mg/kg, Envigo). Additionally, the mice were given 25 mg/kg doxycycline hydrochloride (#D3447, Sigma-Aldrich) resuspended in PBS via intraperitoneal injection once a day for the first two days of the doxycycline regimen to expedite onset of shRNA induction. The tumors were harvested after 1 week of doxycycline treatment. Long-term knockdown was achieved by placing the NSG mice on doxycycline hyclate chow 3 days before orthotopic transplantation and culturing *KPC* GFP-shRNA cells in doxycycline hydrochloride for 3 days before transplant at 2 µg/mL (#D3447, Sigma Aldrich). Transplant recipient NSG mice were maintained on doxycycline hyclate chow until tumors were harvested. The shRNA oligonucleotide sequences are listed below.

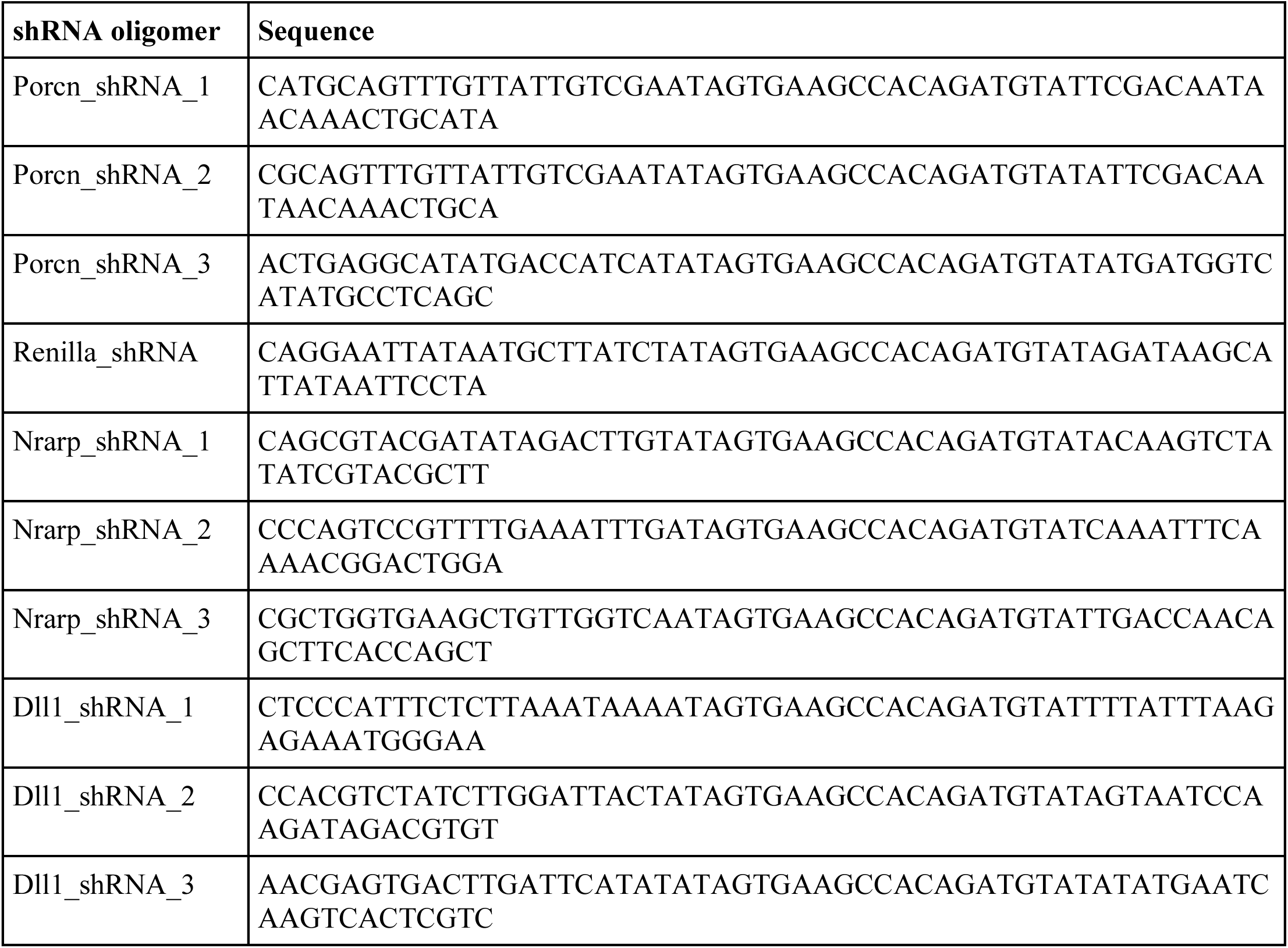

### Gene knockdown in vitro validation

*KPC* PDAC cells expressing the GFP-linked, doxycycline-inducible shRNAs listed above were cultured *in vitro* in doxycycline for 1 week. RNA was extracted and qPCR was performed on the genes of interest. The shRNA for each gene that produced the greatest decrease in expression was used for *in vivo* experiments.

### Gene overexpression in vivo

CRISPR-guided gene activation (CRISPRa) was used to overexpress *Porcn*. We employed the Synergistic Activation Mediator (SAM) CRISPRa system, which is a 3-component system based on (i) a dCas9 fusion to the transcriptional activator VP64 (a tandem repeat of four DALDDFDLDML sequences from *Herpes simplex* viral protein 16, VP16), (ii) a modified gRNA scaffold containing two MS2 RNA aptamers, and (iii) the MS2-P65-HSF1 tripartite synthetic transcriptional activator^53^. In this scenario, sgRNA-dependent recruitment of dCas9-VP64 and MS2-P65-HSF1 to the endogenous *Porcn* locus results in potent transcriptional activation. Ten short guide RNAs (sgRNAs) were screened *in vitro* by qPCR for maximal expression of *Porcn* and the sgRNA providing the highest expression was used for experiments (sgRNA sequence: GGGAGCAGGATTGAG).

### Tumor volume calculation

Volume of *KPC* PDAC orthotopic and subcutaneous tumors was determined by using calipers to measure x-, y-, and z-dimension diameters and using the ellipsoid formula for volume, V = (4/3)*π*(x/2)*(y/2)*(z/2).

### Ablation of WNT-S cells

NSG mice transplanted with *KPC Porcn-DMA* orthotopic tumors were dosed with 50 µg/kg of diphtheria toxin diluted in PBS (#D2189, Millipore Sigma) every other day for 1 week. In PDAC cells harboring the *Porcn-DMA* reporter, DTR is only expressed in *porcupine^+^* (WNT-S) cells.

### CD:LGK974 complexation

CD:LGK974 complexes were prepared by dissolving 100 mg LGK974 and 5656 mg β-cyclodextrin sulfobutyl ether sodium salt (CTD, Inc.) in 20 mL deionized water. The pH was adjusted to 4.0 with 1 N HCl to prepare a clear solution, which was lyophilized into a dry, white powder.

### Pharmacologic porcupine inhibition

To inhibit porcupine enzymatic activity, mice were dosed daily with 10 mg/kg (2.5 µg/µl in a volume of 100 µl) of the small molecule porcupine inhibitor LGK974^17^ in complex with a cyclodextrin carrier (CD:LGK974) by oral gavage. We have previously demonstrated superior solubility and delivery of LGK974 into tumors when complexed with CD^54^.

### In vitro WNT pathway stimulation

To stimulate 2D cell cultures with WNT + R-spondin (WR) media, standard RPMI media was supplemented with 1 µg/ml recombinant mouse R-spondin-1 (SinoBiological) and 100 ng/ml recombinant mouse Wnt3a (#1324-WN, R&D Systems)^19^. Media was changed every 3 days. Alternatively, WNT pathway activation in *KPC* PDAC cells was induced with 2 µM GSK3β inhibitor CHIR99021 (#SML1046, Millipore Sigma) for 12 hours.

### In vitro Notch pathway inhibition

The Notch pathway was inhibited in *KPC* PDAC cells *in vitro* using the γ−secretase inhibitor DAPT ((2S)-N-[(3,5-Difluorophenyl)acetyl]-L-alanyl-2-phenyl]glycine 1,1-dimethylethyl ester) (#D5942, Millipore Sigma). Cells were treated with 20 µM DAPT for 3 days and media was changed daily.

### Isolation of tumor cells

To dissociate tumors into single-cell suspensions, orthotopically transplanted and primary autochthonous *KPCT*; *Lgr5^GFP-IRES-CreER/+^*tumors were finely chopped with razor blades and incubated in a digestion buffer of collagenase IV (#17104019, ThermoFisher Scientific, 0.1 U/ml), Dispase (#354235, Corning, 0.6 U/ml), DNase I (#69182–3; Sigma Aldrich, 10 U/ml), and Soybean Trypsin Inhibitor (#T9003, Sigma Aldrich, 0.1 mg/ml) dissolved in HBSS with Mg^2+^ and Ca^2+^ (#14025076, Thermo Fisher Scientific) in gentleMACS C Tubes (#130-093-237, Miltenyi Biotec) for 42 minutes at 37°C using the gentleMACS Octo Dissociator (#130–096-427, Miltenyi Biotech). After mechanical and enzymatic dissociation, tumor cells were washed with HBSS + Mg^2+^ and Ca^2+^ and filtered through a 100 μm cell strainer and spun at 300 *g* for 5 minutes at room temperature. Cells were then washed with SMEM media and pelleted at 300 *g* for 5 minutes at 4°C. The supernatant was removed, and the pellet resuspended in fluorescence-activated cell sorting (FACS) buffer (200 mM EDTA with 2% of heat-inactivated FBS in PBS) before passing through a 40 μm strainer. To stain cells for FACS, Fc receptor was blocked at room temperature for 5 minutes with rat anti-mouse CD16/CD32 (Mouse BD Fc Block, #553142, BD Biosciences) in FACS buffer and incubated for 20 minutes with a mix of 5 PE-Cy7 conjugated antibodies binding CD45 (#25-0451-82, Invitrogen, 1:160), CD31 (#25-0311-82, Invitrogen, 1:40), CD11b (#25-0112-81, Invitrogen, 1:160), TER-119 (#116222, Biolegend, 1:80), and F4/80 (#5-4801-82, Invitrogen, 1:40), referred to as stroma. Either 300 nM of DAPI (#D9542, Sigma Aldrich, 5 μg/ml) or ZombieNIR (#423106, Biolegend, 1:1000) was used as a live/dead marker. To separate WT cells from Porcn-CRISPRa or Porcn-shRNA cells in mixed population tumor transplants, the samples were additionally stained with a PerCP-Cy5.5 conjugated antibody binding Thy1.1 (#202515, Biolegend, 1:160). Cell sorting was carried out on a BD FACSAria sorter (BD Biosciences) for live cells. Cells were sorted into SMEM with 2% of heat inactivated FBS.

### Cell culture and reagents

Primary 2D mouse cell lines were generated from autochthonous *KP* and *KPT* PDAC tumors by dissociation and FACS, as described above, and cultured in RPMI (#11875119, Gibco) supplemented with 1% GlutaMax (#35050061, Gibco), 1% Penicillin/Streptomycin (Pen/Strep) (#15070063, Gibco), and 2% Heat-Inactivated FBS. After 24 h, media was changed to remove non-adherent cells and replaced by Advanced DMEM/F12 (#12634028, Gibco) with 1% GlutaMax (#35050061, Gibco), 1% Pen/Strep (#15070063, Gibco), and 10% FBS. Cells were passaged using 0.25% Trypsin-EDTA (#25200056, Gibco).

### Tumor sphere culture

FACS-purified mScarlet^+^, 7TCF-eGFP^+^, and mScarlet^−^/7TCF-eGFP^−^ cells isolated from orthotopic *7TCF/Porcn-DMA* reporter PDAC tumors as described above were resuspended in tumor sphere culture medium [Advanced DMEM/F12 (#12634028, Gibco), 2% FBS (#SH30910.03, Hyclone), 1% GlutaMax (#35050061, Gibco), 1% Pen/Strep (#15070063, Gibco)] and mixed with 50 µl Matrigel (#CB-40230C Fisher Scientific). The cell-Matrigel mix was placed in 24-well plates (#353047, Thermo Fisher Scientific) and incubated at +37°C for 30 min. Either 1000 cells of one population or 500 cells each of two populations were plated together. Tumor sphere culture media (750 µl) was added and replaced every 3 days during culture. Tumor spheres were imaged with an EVOS M5000 microscope (ThermoFisher) and quantified manually.

### Measurement of WNT secretion capacity

*KPC* PDAC cells harboring the *Porcn-DMA* reporter were plated in white-walled 96 well plates (#CLS3922, Corning) at 10,000 cells per well. After 24 hours, the cells were transfected with a mammalian cDNA expression plasmid enabling constitutive expression of human WNT3a. The supernatant from these cells was collected after 48 hours, filtered through a .45 µm filter, and transferred to human cells expressing the TOPFLASH Firefly (M50) 7TCF luciferase reporter (Addgene plasmid #12456)^28^ and the constitutive Renilla luciferase pRL-SV40P (Addgene plasmid #27163; TOPFLASH assay)^27^. Luminescence was measured using the Dual-Reporter Assay (#E1910, Promega) per the manufacturer’s instructions with an BioTek Cytation 5 (Agilent). 7TCF-Firefly signal was normalized to Renilla luminescence in each well.

### Generation of DTR-mScarlet-AkaLuc-Porcn (Porcn-DMA) reporter cell lines

Homology arms 500 bp in length 5’ and 3’ from the start codon of *Porcn* exon 1 were amplified from C57/BL6 genomic DNA using high-fidelity PCR (#M0494, NEB Q5 polymerase). A homology-directed repair template donor vector where *frt-TAG-bGlobinpA-(PGK-Hygro-pA)i-frt-DTR-P2A-mScarlet-AkaLuc-T2A* is flanked by the homology arms was cloned into the pUC19 plasmid backbone using Gibson Assembly (#E2611, NEB). A sgRNA (CCATCTGTGTGGGTCCACAA) targeting the beginning of exon 1 of *Porcn* was cloned by ligation into a mammalian expression vector also expressing Cas9 and GFP (pX458)^55^. *KPC* PDAC cell lines were co-transfected with the *Porcn-DMA* donor vector and sgPorcn-pX458, followed by antibiotic selection with hygromycin. Single cell clones were picked from the pool of targeted cells and expanded. Correct targeting of clones was validated by genotyping using primers specific to DTR, mScarlet, AkaLuc, and the *Porcn* genomic locus across the homology arms. The oligonucleotides used for the genotyping of *Porcn-DMA* are listed below.

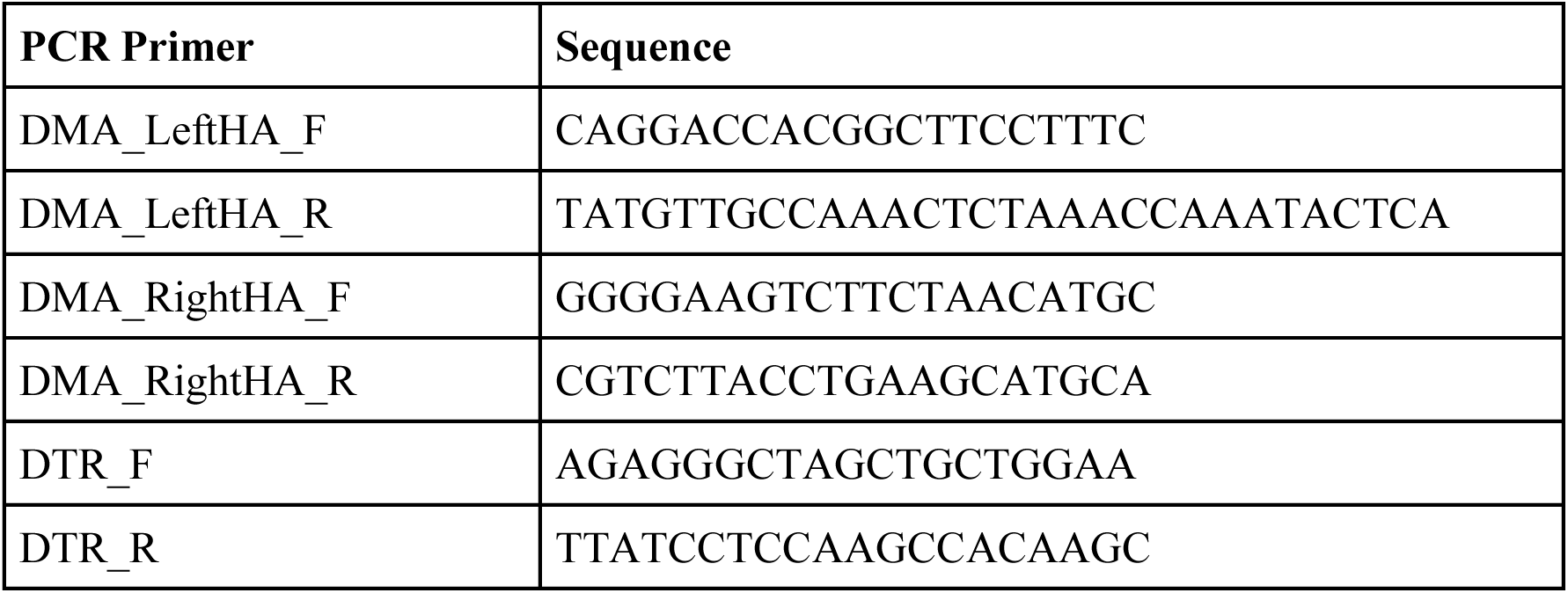
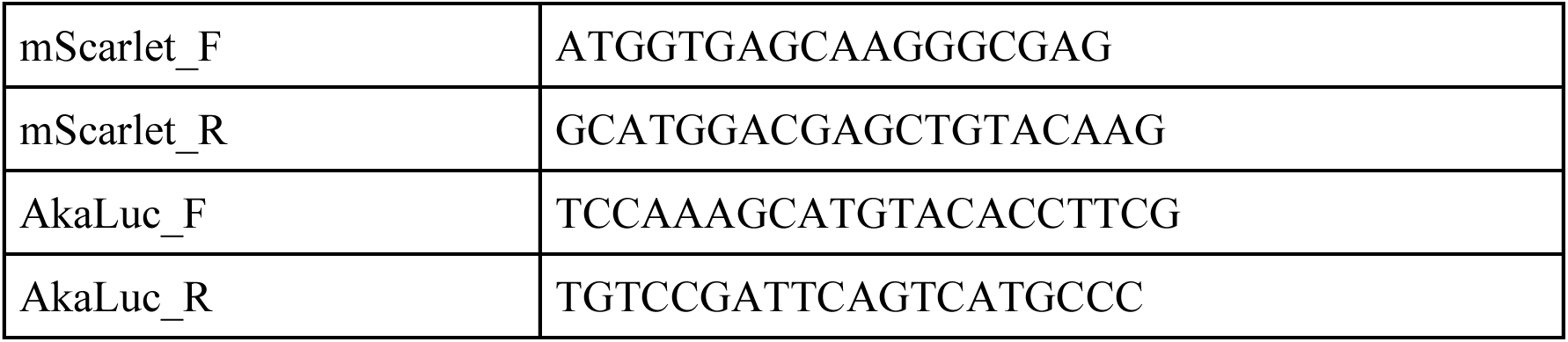

The clones with a correctly targeted *Porcn-DMA* reporter were then transduced with adenoviral FlpO (University of Iowa, Viral Vector Core) to remove the *frt-TAG-bGlobinpA-(PGK-Hygro-pA)i-frt* “STOP” cassette. Complete excision of the STOP cassette was validated by genotyping using primers specifically amplifying across DTR and the left homology arm. Clones with the STOP cassette removed were FACS-isolated into single-cell clones by mScarlet fluorescent signal and subsequently expanded in culture.

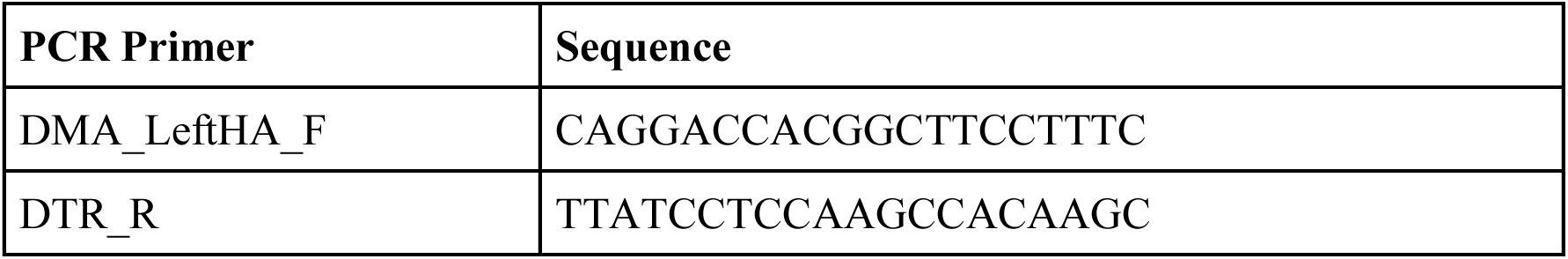

### Generation of Porcn-DTR-mScarlet-AkaLuc-CreER (Porcn-DMAC) reporter cell line

Homology arms ∼900 bp in length 5’ and 3’ to the end of *Porcn* exon 14 were amplified from C57/BL6 genomic DNA using high-fidelity PCR (#M0494, NEB Q5 polymerase). A homology-directed repair template donor vector where *frt-PGK-Hygro-pA-frt-IRES2-DTR-P2A-HAtag-mScarlet-AkaLuc-T2A-CreER* is flanked by the homology arms was cloned into the pUC19 plasmid backbone as described above. A sgRNA (ATATGCCTCAGCCTATGAGA) targeting the end of exon 14 of *Porcn* was cloned by ligation into a mammalian expression vector also expressing Cas9 and GFP (pX458)^55^. Targeting, selection, single-cell cloning and genotyping of *Porcn-DMAC* cells was done as described above. The additional oligonucleotides used for the genotyping of *Porcn-DMAC* are listed below.

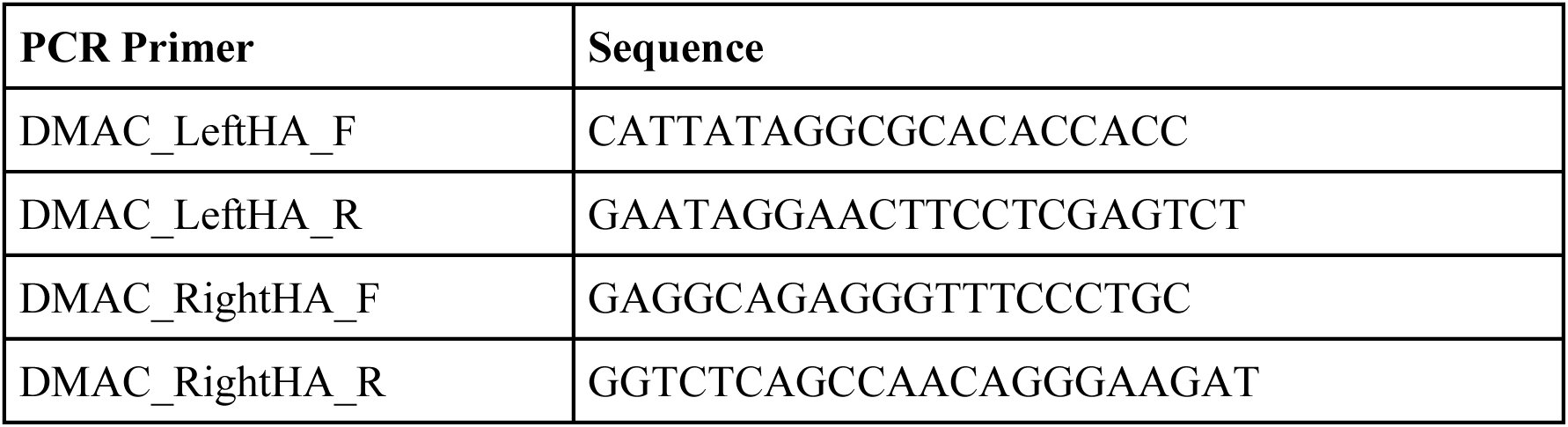

The clones with a correctly targeted *Porcn-DMAC* reporter were then transduced with adenoviral FlpO (University of Iowa, Viral Vector Core) to remove the *frt-PGK-Hygro-pA-frt* “STOP” cassette. Complete excision of the STOP cassette was validated by genotyping using primers specifically amplifying across IRES2 and the left homology arm. Clones with the STOP cassette removed were FACS-isolated into single-cell clones by mScarlet fluorescent signal and subsequently expanded in culture.

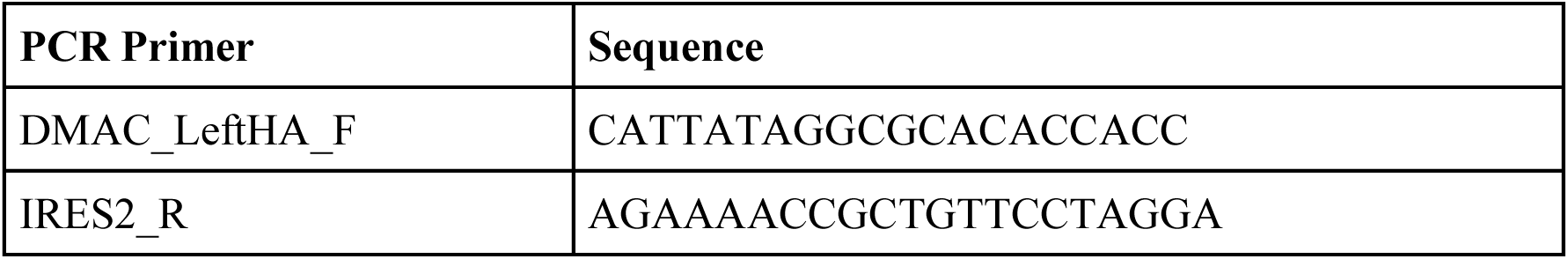

### Generation of 7TCF (WNT-R) reporter cell line and WNT-S or WNT-R lineage tracing cell lines

To generate the WNT-R reporter system, a WNT-responsive *7TCF-eGFP* or *7TCF-tagBFP* cassette was cloned upstream of a constitutively expressed *PGK-GaussiaLuciferase-P2A-iRFP670* cassette into a lentiviral plasmid backbone using Gibson Assembly (#E2611, NEB). Lentivirus was generated in HEK293 cells and used to transduce *KPC* PDAC *Porcn-DMA* reporter cells *in vitro*. Transduced cells were isolated by FACS based on iRFP670 expression. To create the cell lines enabling lineage-tracing of the WNT-S or WNT-R states, the *7TCF-eGFP-PGK-GaussiaLuciferase-P2A-iRFP670* reporter was modified to include *EFS-FLEx-tagBFP2*. FLEx (flip-excision) constructs contain two pairs of heterologous loxP sites flanking the “flexed” construct (here, *tagBFP2*). When exposed to Cre, the “flexed” construct is inverted, followed by an excision of the loxP sites that locks the construct in the flipped orientation. With *EFS-FLEx-tagBFP2*, after Cre recombination the constitutive promoter elongation factor 1a short (EFS) drives expression of tagBFP2. Lentivirus was generated as described above and used to transduce *KPC* PDAC *Porcn-DMA* reporter cells *in vitro* for WNT-R lineage-tracing and *DMAC-Porcn* reporter cells for WNT-S lineage-tracing.

### Human PDAC tumor tissue microarrays

Formalin-fixed and paraffin-embedded (FFPE) surgical tissue samples (n=261) were collected from the archives of the Department of Pathology, Helsinki University Hospital. The diagnosis of PDAC was confirmed by an experienced pathologist. The most representative tumor regions were marked on hematoxylin and eosin-stained slides. Multipunch Tissue Microarray blocks (TMAs) were prepared from the marked areas of the tissue samples. Six 1 mm cores were cut from different areas of each sample with a semiautomatic tissue microarrayer (Tissue Arrayer 1, Beecher Instruments Inc.). TMA blocks were cut into 4 µm sections that were mounted on positively charged slides for immunohistochemistry. The study was approved by the Surgical Ethics Committee of Helsinki University Hospital and carried out in accordance with the Declaration of Helsinki. Written informed consent was obtained from all participants prior to sample collection.

### Immunohistochemistry

Slides were baked for 30 minutes at 60°C, deparaffinized, rehydrated with an alcohol series, then washed with PBS. Antigen retrieval was performed for 20 minutes at 95°C using Tris-EDTA antigen retrieval buffer. Slides were left to cool before being washed with PBS-T. Bloxall was added to each section and slides were incubated for 10 minutes at RT. Primary antibody (PORCN, #ab201793, AbCam; or GFP, #2956, Cell Signaling Systems) was diluted in TNB buffer (PerkinElmer). Slides were incubated at 4°C overnight. The following day, slides were washed with PBS-T and secondary antibody. Slides were incubated for 1 hour at RT, then washed with PBS-T. DAB was added to each section and sections were incubated until brown. PBS-T was used to stop the reaction. Counterstaining was performed using hematoxylin and bluing solution. Slides were dehydrated by moving them through the reverse rehydration sequence and mounted with toluene, then left to dry. Images were acquired using a Mirax Midi-Scanner (Carl Zeiss AG). Samples were evaluated in CaseViewer (3DHISTECH) and were scored according to staining intensity as follows: 0 (no staining), 1 (weak staining), 2 (moderate staining), and 3 (strong staining). Scoring was performed blinded to sample ID.

### Tissue histology and immunofluorescence

For immunofluorescence of cryosections, tumors harvested at experimental endpoints were fixed in 10% neutral buffered formalin (#HT501128, Millipore Sigma), embedded in OCT (#23-730-571, Fisher Scientific), and cut into 10 µm sections. Sections were placed on SuperFrost microscope slides (#12-550-15, Fischer Scientific) and used for staining. Sections were fixed in acetone for 10 minutes, washed in PBS, and blocked for 30 minutes in PBS with 0.3% Triton-X, 0.2% BSA, and 5% donkey serum (#D9663, Sigma-Aldrich). The slides were incubated with primary antibodies overnight at 4°C in blocking buffer. The following primary antibodies were used: GFP (#ab5450, Abcam, 1:500), porcupine (#MABS21, Millipore Sigma, 1:500), FLAG (#ab205606, Abcam, 1:500), RFP (#600-401-379, Rockland, 1:500), NRARP (#BS-11914R, Thermo Fisher Scientific, 1:500), DLL1 (#AF5026, R&D Systems, 1:40), and Ki67 (#14-5698-82, Thermo Fisher Scientific, 1:100). After washing and permeabilizing with 0.1% PBS-TritonX 100, the slides were incubated with secondary antibodies (#A-11055, #A-11015, #A-21206, #A-11037, #A-31573, #A-21448, Thermo Fisher Scientific, 1:500) for 2 hours at room temperature. The tissues were counterstained with DAPI (#D9542, Sigma Aldrich, 5 μg/ml) for 10 minutes and mounted on cover slips with Mowiol mounting reagent (#475904, Millipore Sigma). Images were acquired on a Zeiss Axio Imager Z2 with ZEN 2.3 software using 20x or 40x objectives or a Mirax Midi-Scanner (Carl Zeiss AG).

For immunofluorescence of formalin-fixed paraffin-embedded (FFPE) tissues, tumors harvested at experimental endpoints were fixed in 10% neutral buffered formalin (#HT501128, Millipore Sigma), embedded in paraffin, and cut into 5 µm sections. The slides were heated for 30 minutes at 60°C, deparaffinized, rehydrated with an alcohol series and incubated for 20 minutes in a pressure cooker on 95°C in Tris-EDTA antigen retrieval buffer (#E1161, Sigma-Aldrich). The tissues were subsequently washed in PBS, permeabilized in 0.3% PBS-Triton X-100 for 40 minutes and blocked in PBS with 0.1% Triton X-100, 2% BSA, and 5% donkey serum (#D9663, Sigma-Aldrich). After PBS wash, the slides were incubated for 1 hour at room temperature in secondary antibodies (see above). The tissues were counterstained with DAPI (#D9542, Sigma Aldrich, 5 μg/ml) for 10 minutes and mounted on cover slips with Mowiol mounting reagent (#475904, Millipore Sigma). Images were acquired on a Zeiss Axio Imager Z2 with ZEN 2.3 software using 20x or 40x objectives.

### In Situ Hybridization

Single-molecule mRNA *in situ* hybridization was performed on FFPE tissues using the Advanced Cell Diagnostics RNAscope 2.5 HD Detection Kit (#322360, ACD) and probes Hs-PORCN (#450741, ACD), Hs-PORCN-C2 (#450741-C2, ACD), Hs-LGR5-C2 (#311021-C2, ACD), Hs-AXIN2-C2 (#400241-C2, ACD), Hs-NRARP (#311871, ACD), Hs-DLL1 (#532631, ACD), according to the manufacturer’s instructions.

### Quantitative PCR (qPCR)

RNA was isolated from whole tumors or cell populations isolated from primary tumors using the RNeasy Plus Micro kit (#74034, Qiagen) per the manufacturer’s instructions. RNA was isolated from 2D cell culture populations using the RNeasy Plus Mini kit (#74134, Qiagen) per the manufacturer’s instructions. cDNA was synthesized with the PrimeScript RT Reagent kit (#RR037B, Takara). qPCR was performed in technical quadruplicates with 1-2 μl of cDNA (diluted 1:10 if necessary) using the PowerUP SYBR mix (#A25778, Applied Biosystems) and analyzed on the QuantStudio 7 Flex Real-Time PCR System. The DDCT method was used to quantify relative gene expression, normalized to *ActinB*. The oligonucleotides used for qPCR amplification in this study are listed below.

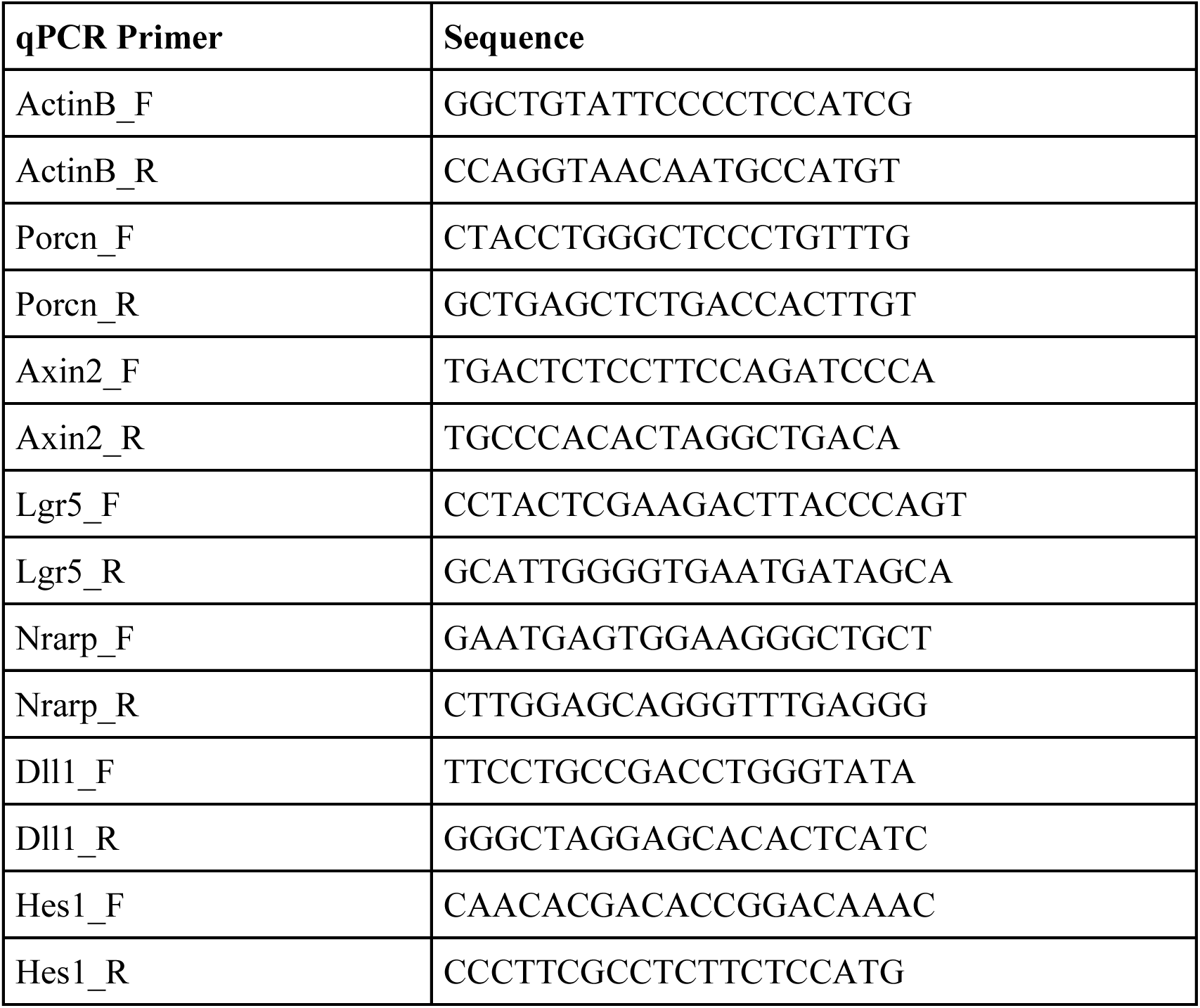

### Computational analyses

#### Bulk RNA sequencing analysis

WNT-R, WNT-S, WNT-R/S, and DN cells from *KPC 7TCF-eGFP*/*Porcn-DMA* PDAC orthotopic tumors were FACS-isolated, RNA was extracted, and libraries were prepared for SMARTerSeq2, *n* = 4 mice/group. FASTQ files were processed using the standard DESeq2 pipeline (version 1.24.0). Reads were aligned to the GRCm38 mouse genome. Count matrices were generated, and differential gene expression analysis was performed using DESeq2 (**Table S1**). Data were filtered to only include genes with 10 or more counts in 2 or more samples. The significance of differential expression between WNT-R, WNT-S, WNT-R/S, and DN populations was quantified by one-way ANOVA.

#### Single cell RNA sequencing

Single-cell data from *KPCT* GEMM PDAC tumors was generated in our prior study, with processing, quality control, clustering, and embedding as previously described^9^. Data analysis was performed in Python and R using a combination of published packages and custom scripts centered around the *scanpy* analysis package ^56^. To de-noise and recover missing gene values, MAGIC imputation was applied to the normalized count matrix using default parameters^57^. Pseudotemporal orderings of tumor cells were generated using the diffusion pseudotime function implemented in *scanpy*^58^. For heatmap visualizations, the mean expression of a gene of interest was plotted over pseudotime using a sliding window covering 100 cells.

#### PORCN expression in human PDAC patients

Bulk RNA-seq from GTEx (Genotype-Tissue Expression Program, gtexportal.org) and TCGA (The Cancer Genome Atlas, portal.gdc.cancer.gov) data were obtained from the UCSC Toil RNA-seq recompute compendium (xena.ucsc.edu), a uniformly re-aligned and re-called expression dataset that enables cross-program comparisons free of computational batch-effects^59^. TCGA tumors were limited to the set of 150 samples confidently identified as PDAC resulting in sample counts as follows: GTEx normal samples *n* = 165, TCGA tumor samples *n* = 150. Normalized gene expression counts for *PORCN* were standardized and visualized using an empirical cumulative density function plot. The significance of the difference between tumor and normal tissue expression was quantified using a Kolmogorov-Smirnov test.

## Acknowledgments

We thank members of the Tammela laboratory and A. Smogorzewska for helpful discussions; M. Sherman for comments on the manuscript; D. Pe’er for insightful feedback and help in funding acquisition; A.J. Bhutkar for analysis of *PORCN* expression in human patient gene expression data; E. di Stanchina for mouse surgeries; R. Gardner for FACS support; K. Manova and M. Tipping for histology support; N. Mohibullah for next-generation sequencing; H. Alcorn for laboratory management; and K. Bastl, J. Chan, S. Ding, M. Gregory, G. Hartmann, A. Hudson, M. Kim, K. Krause, M. Raghavan, H. Styers, and C. Sussman for help with experiments. This work was supported by the NIH/NCI (R37-CA244911) and Cancer Center Support Grant (P30-CA08748 to MSKCC), the Starr Cancer Consortium, and The Robertson Foundation (via the Josie Robertson Investigators Program at MSKCC). S.T. was supported by a fellowship from NIH/NCI (F30-CA254120) and a Medical Scientist Training Program grant from NIGMS/NIH under award number T32GM007739 to the Weill Cornell/Rockefeller/Sloan Kettering Tri-Institutional MD-PhD Program. T.T. was supported by Scholarships from the American Cancer Society, the Rita Allen Foundation, and the V Foundation. We acknowledge the use of the Integrated Genomics Operation, Antitumor Assessment, Flow Cytometry, and Histology Core Facilities at Sloan Kettering Institute, funded by CCSG P30-CA08748.

## Author contributions

S.R.T. and T.T. conceived and designed the study. S.R.T., O.G.H., M.H., K.W., K.L.P., and F.S.R. performed experiments and analyzed data. S.R.T., A.S., and Y.J.H. performed computational analysis. X.H. and M.J.M. formulated cyclodextrin-LGK974 complexes. A. L. performed the mathematical modeling. C.H. provided human pancreas cancer tissue microarrays. S.W.L., L.E.D., and F.S.R. contributed gene perturbation expertise and reagents. S.R.T. and T.T. wrote the manuscript. All authors provided input and reviewed the manuscript. T.T. supervised the study.

## Declaration of interests

**O.G.H.:** Employee of Loxo @Lilly with ownership interest (including stocks). **S.W.L.:** Founder and member of the scientific advisory board of Blueprint Medicines, Mirimus, ORIC Pharmaceuticals, and Faeth Therapeutics; advisory role and equity interests in Constellation Pharmaceuticals and PMV Pharmaceuticals. **L.E.D.:** Advisory role and equity interests in Mirimus Inc. **T.T.:** Advisory role and equity interests in Lime Therapeutics; the Tammela Lab receives research support from Ono Pharmaceuticals Co., Ltd (unrelated to this work).

---

**Figure S1.**
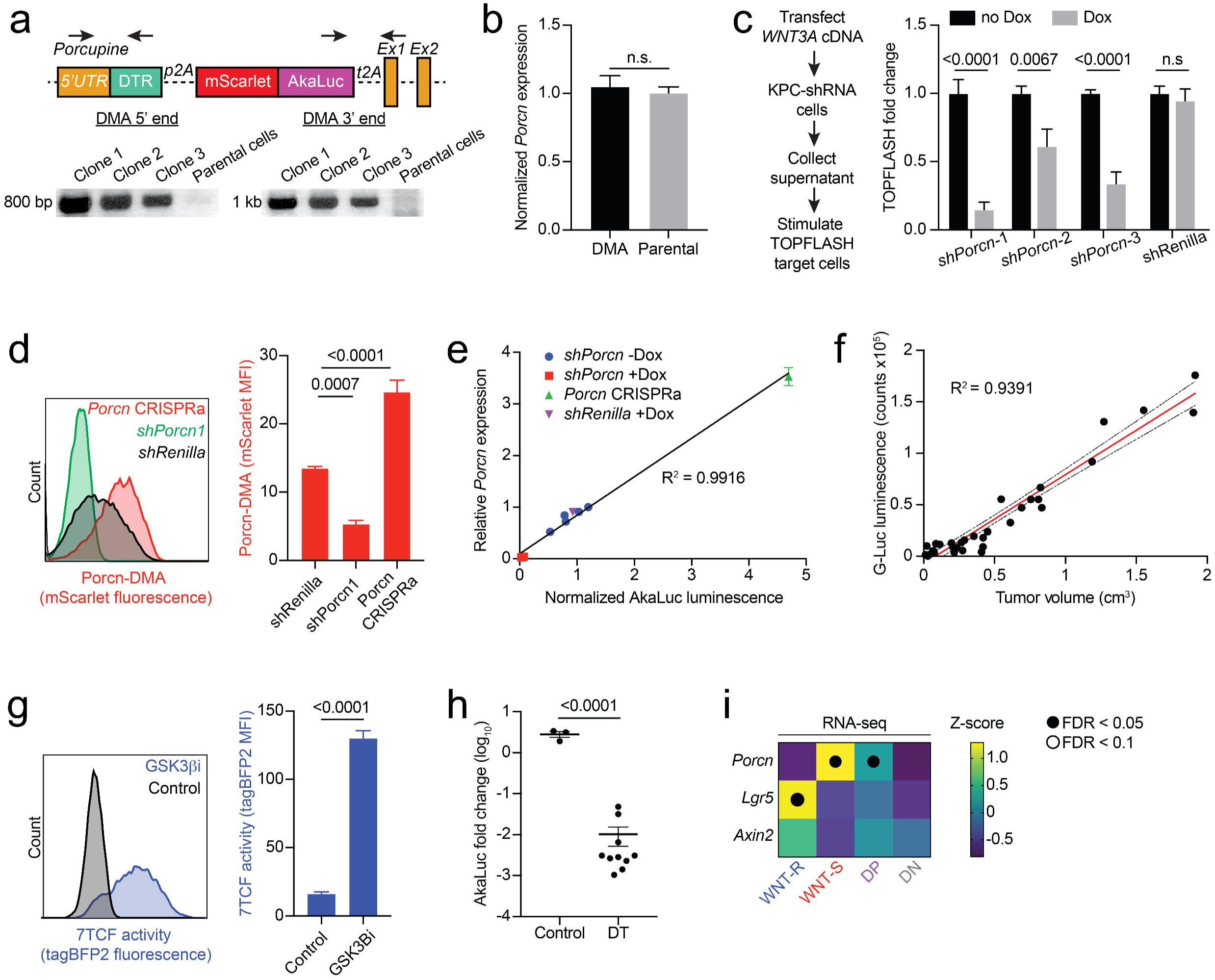
Construction and validation of *7TCF/Porcn-DMA* reporter cell lines, related to Figure 1. (**a**) Schematic of the *Porcn-DMA* reporter (top) and genotyping PCRs for integration in *KPC* PDAC cells (bottom). Black arrows indicate the binding sites of genotyping primers. (**b**) Quantitative PCR (qPCR) for *Porcn* mRNA in cultured *KPC* cells targeted with *Porcn-DMA* vs. parental *KPC* cells; *n* = 4-12 technical replicates. (**c**) Normalized WNT secretion capacity of *Porcn-DMA KPC* PDAC cells *in vitro*. *Porcn-DMA KPC* PDAC cells were transfected with a plasmid encoding *WNT3A* and supernatant was used to stimulate 293T cells transfected with a WNT pathway reporter assay (TOPFLASH assay); *n* = 9 technical replicates. (**d**) Representative flow cytometry histogram of mScarlet expression in *Porcn-DMA* reporter PDAC cells lentivirally transduced with *Porcn* shRNA or CRISPRa *in vitro* (left); quantification of mScarlet mean fluorescence intensity (MFI, right). (**e**) Comparison of *Porcn* expression by qPCR and AkaLuc bioluminescence in *Porcn-DMA* reporter PDAC cells lentivirally transduced with *Porcn* shRNA or CRISPRa *in vitro*. (**f**) Comparison of *Gaussia* luciferase (G-Luc) signal and tumor volume in NSG mice with subcutaneous *7TCF/Porcn-DMA* reporter PDAC allografts. Red line indicates linear regression; black dotted lines indicate 95% confidence interval. (**g**) Representative flow cytometry histogram of 7TCF-tagBFP2 expression in *7TCF/Porcn-DMA* reporter PDAC cells subjected to GSK3β inhibitor CHIR99021 or vehicle control *in vitro* (left); quantification of 7TCF-tagBFP2 MFI (right). (**h**) Quantification of AkaLuc luminescence in mice bearing orthotopically transplanted *7TCF/Porcn-DMA* reporter PDAC tumors after 7 days of diphtheria toxin (DT) administration. (**i**) RNA-seq analysis of *Porcn, Lgr5,* and *Axin2* expression in primary WNT-R, WNT-S, WNT-R/S, and DN cells isolated from orthotopic PDAC *7TCF/Porcn-DMA* reporter allografts. Expression is quantified as a row-normalized z-score; *n* = 4 mice. Unpaired *t* tests were used to test significance in (**b**), (**g**) and (**h**). Two-way ANOVA was used in (**c**) and (**d**) to test for statistical significance. One-way ANOVA with FDR was used in (**i**) to test for statistical significance. Open circles indicate FDR < 0.1; closed circles indicate FDR < 0.05. Error bars indicate SEM.

**Figure S2.**
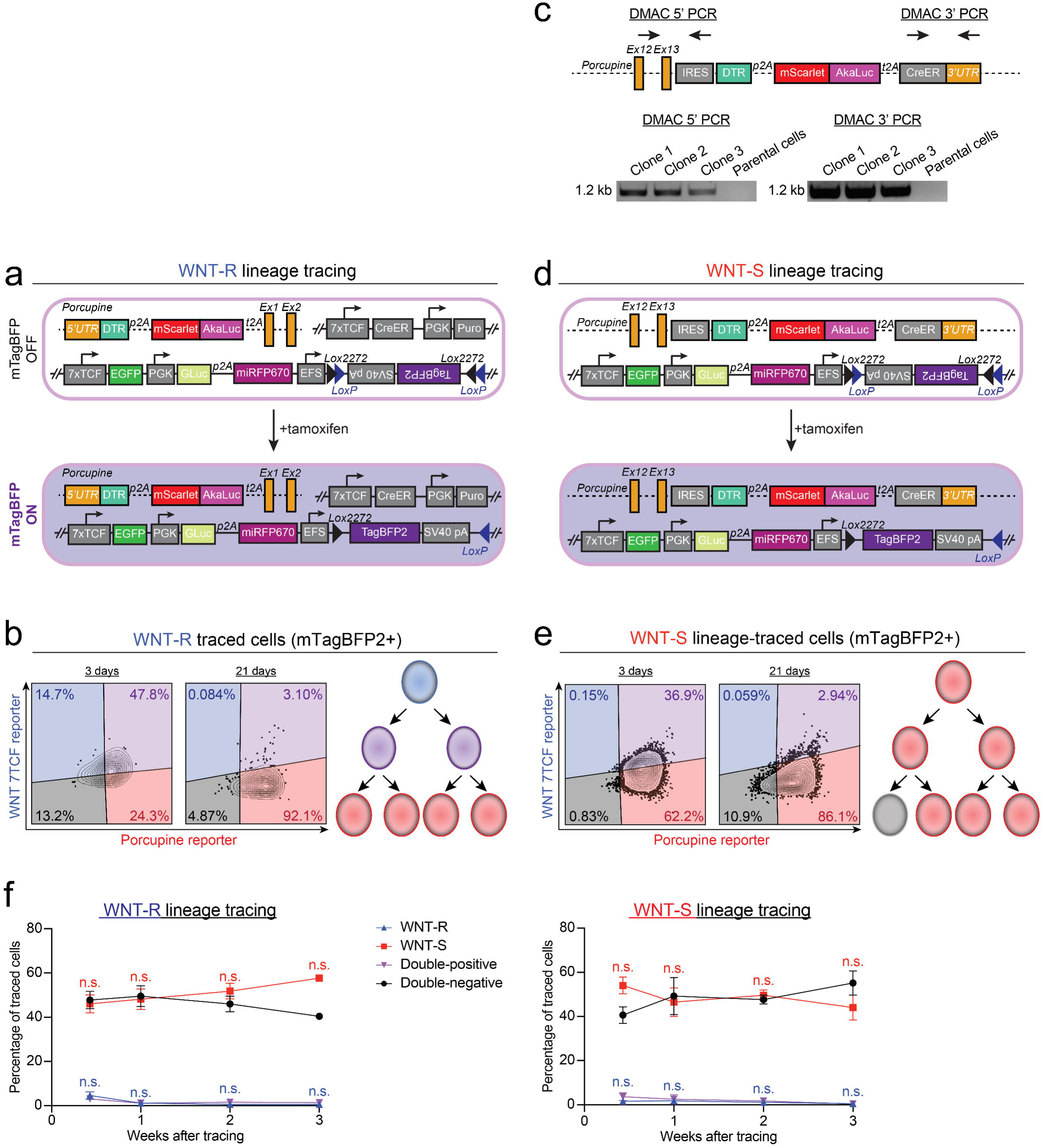
Genetic systems and outcome of lineage-tracing WNT-R and WNT-S cells in primary PDAC tumors, related to Figure 1. (**a**) Genetic systems used for WNT-R lineage tracing in *7TCF-eGFP-CreER/Porcn-DMA KPC* cell lines. These cells contain a *7TCF-eGFP* reporter that also contains a Cre-responsive tagBFP2 that becomes active upon Cre recombinase activation. A second lentiviral construct containing *7TCF-CreER* enables lineage tracing of 7TCF^+^ (WNT-R) cells following a single pulse of tamoxifen (bottom). (**b**) Representative flow cytometry plots of orthotopic *7TCF-eGFP-CreER/Porcn-DMA KPC* WNT-R lineage tracing allografts 3 days and 14 days after tamoxifen dosing (left). Note that only the traced, tagBFP2^+^ cells are shown in the flow cytometry plots. Diagram summarizing WNT-R lineage tracing results (right): The WNT-R (blue) and the WNT-R/S (purple; double-positive, DP) are initially labeled, and eventually give rise to WNT-S cells (red). The DP and the WNT-S cells expand by proliferation. (**c**) Genetic system for visualizing and lineage tracing the WNT-S population engineered into *KPC* cell lines. In addition to the components of *Porcn-DMA*, *Porcn-DMAC* contains **<u>C</u>**reER for lineage tracing (top). Genotyping PCRs for integration of *Porcn-DMAC* in *KPC* PDAC cells (bottom). Black arrows indicate the binding sites of genotyping primers. (**d**) Genetic systems for WNT-S lineage tracing *7TCF-eGFP/Porcn-DMAC KPC* cell lines. These cells contain a *7TCF-eGFP* reporter that also contains a Cre-responsive tagBFP2 that becomes active upon Cre recombinase activation. CreER placed under the control of endogenous *Porcn* regulatory elements enables lineage tracing of porcupine^+^ (WNT-S) cells following a pulse of tamoxifen (bottom). (**e**) Representative flow cytometry plots of orthotopic *KPC 7TCF-eGFP/Porcn-DMAC* WNT-S lineage tracing allografts 3 days and 14 days after tamoxifen administration (left). Note that only the traced, tagBFP2^+^ cells are shown in the flow cytometry plots. Diagram summarizing WNT-S lineage tracing results (right): The DP and WNT-S cells are labeled, followed by rapid differentiation of the DP cells into the WNT-S state, which expand by proliferating rapidly. A small subset of the traced cells become double-negative (DN). (**f**) Quantification of WNT-R, WNT-S, WNT-R/S, and DN populations in the non-lineage-traced (tagBFP2^−^) fraction of 7*TCF-eGFP-CreER/Porcn-DMA* reporter PDAC allografts (WNT-R lineage tracing experiment, left) or *7TCF-eGFP/Porcn-DMAC* reporter PDAC allografts (WNT-S lineage tracing, right). Unpaired *t* tests were used to test significance compared to baseline at each time point in (**f**). Error bars indicate SEM.

**Figure S3.**
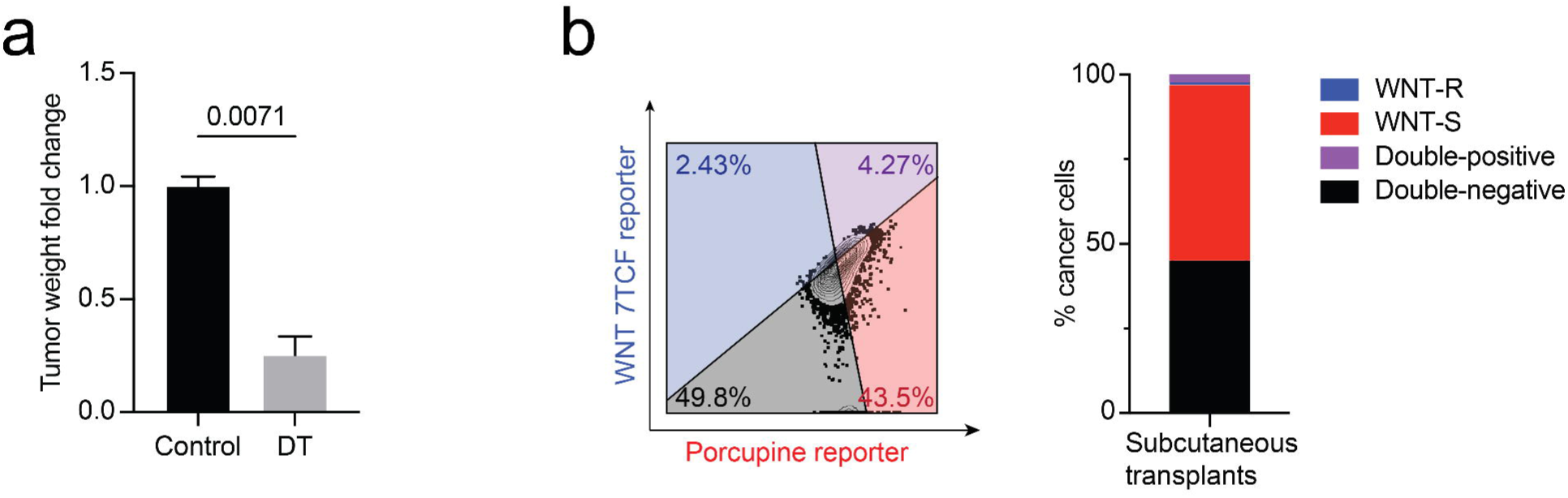
WNT-S state is required for growth of primary and metastatic PDAC, related to Figure 2. (**a**) Weight of orthotopic *7TCF/Porcn-DMA* reporter PDAC allografts harvested after 1 week of DT treatment; *n* = 2-3 mice/group. (**b**) Representative flow cytometry plot from *7TCF/Porcn-DMA* reporter PDAC tumors in a subcutaneous transplant model (left). Stacked bar graph showing percentage of WNT-S, WNT-R, WNT-R/S, and DN cells in tumors (right); *n* = 6 mice/group. Unpaired *t* test was used to test significance in (**a**). Error bars indicate SEM.

**Figure S4.**
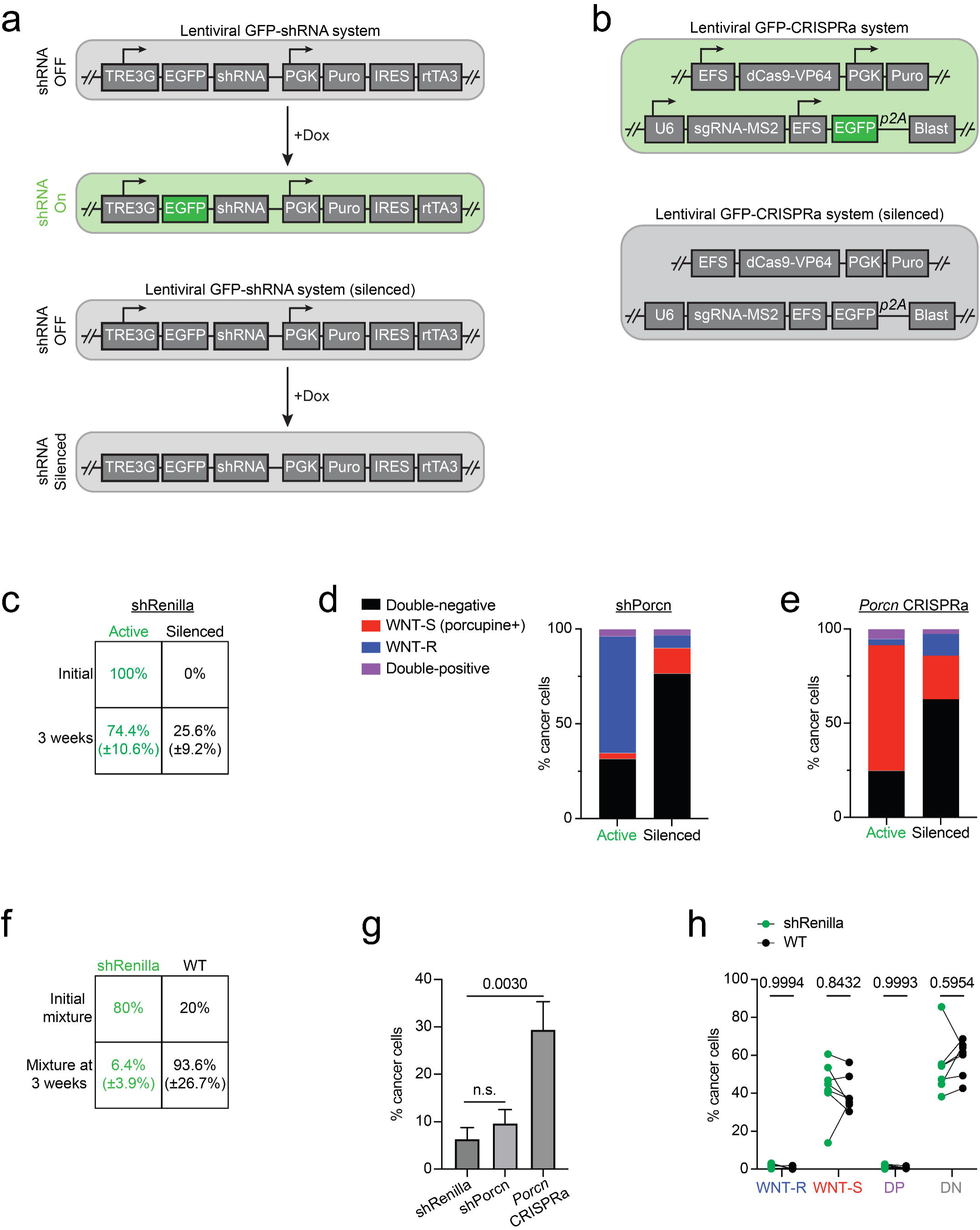
Genetic systems of *Porcn* loss- and gain-of-function reveal WNT-S/WNT-R equilibrium is essential for PDAC growth, related to Figure 3. (**a**) Lentiviral constructs used for doxycycline (Dox)-inducible shRNA knockdown. Binding of Dox to reverse tetracycline-controlled transactivator-3 (rtTA3) enables rtTA3 to activate a 3^rd^-generation tetracycline response element (TRE3G) promoter and drive expression of GFP and the linked shRNA. Loss of GFP in the presence of Dox indicates that the construct has been silenced (bottom). (**b**) Lentiviral constructs used for CRISPR-guided gene activation (CRISPRa, see STAR Methods). Loss of GFP indicates that the construct has been silenced (bottom). (**c**) Quantification of cells in orthotopic *7TCF/Porcn-DMA* reporter allografts harboring *Renilla* shRNA-GFP with the genetic system active (GFP+) or silenced (GFP-) initially and at tumor harvest. (**d**, **e**) Stacked bar graphs showing the percentage of WNT-S, WNT-R, WNT-R/S, and DN cells in orthotopic allografts of *7TCF/Porcn-DMA* reporter PDAC cells harboring *Porcn* shRNA-GFP (**d**) or *Porcn* CRISPRa-GFP (**e**); *n* = 5-7 mice/group. (**f**) Quantification of *Renilla* shRNA-GFP^+^ vs. wild-type cells in orthotopic mixed *7TCF/Porcn-DMA* reporter allografts at transplantation (Initial mixture) and at harvest. (**g**) Quantification of *Porcn* shRNA-GFP^+^, *Porcn* CRISPRa-GFP^+^, and *Renilla* shRNA-GFP^+^ cells from orthotopic mixed *7TCF/Porcn-DMA* reporter allografts at the time of harvest, *n* = 8-12 mice/group. (**h**) Quantification of cell states in experiment described in (**f**). PDAC subpopulations harboring GFP-linked shRenilla (green dots) or wild type cells (black dots) in the same tumor are connected by a line; *n* = 7 mice/group. Chi-squared test was used in (**c**) and (**f**) to test for statistical significance. Two-way ANOVA was used in (**g**) and (**h**) to test for statistical significance. Values in parentheses in (**c**) and (**f**) indicate SD. Error bars indicate SEM.

**Figure S5.**
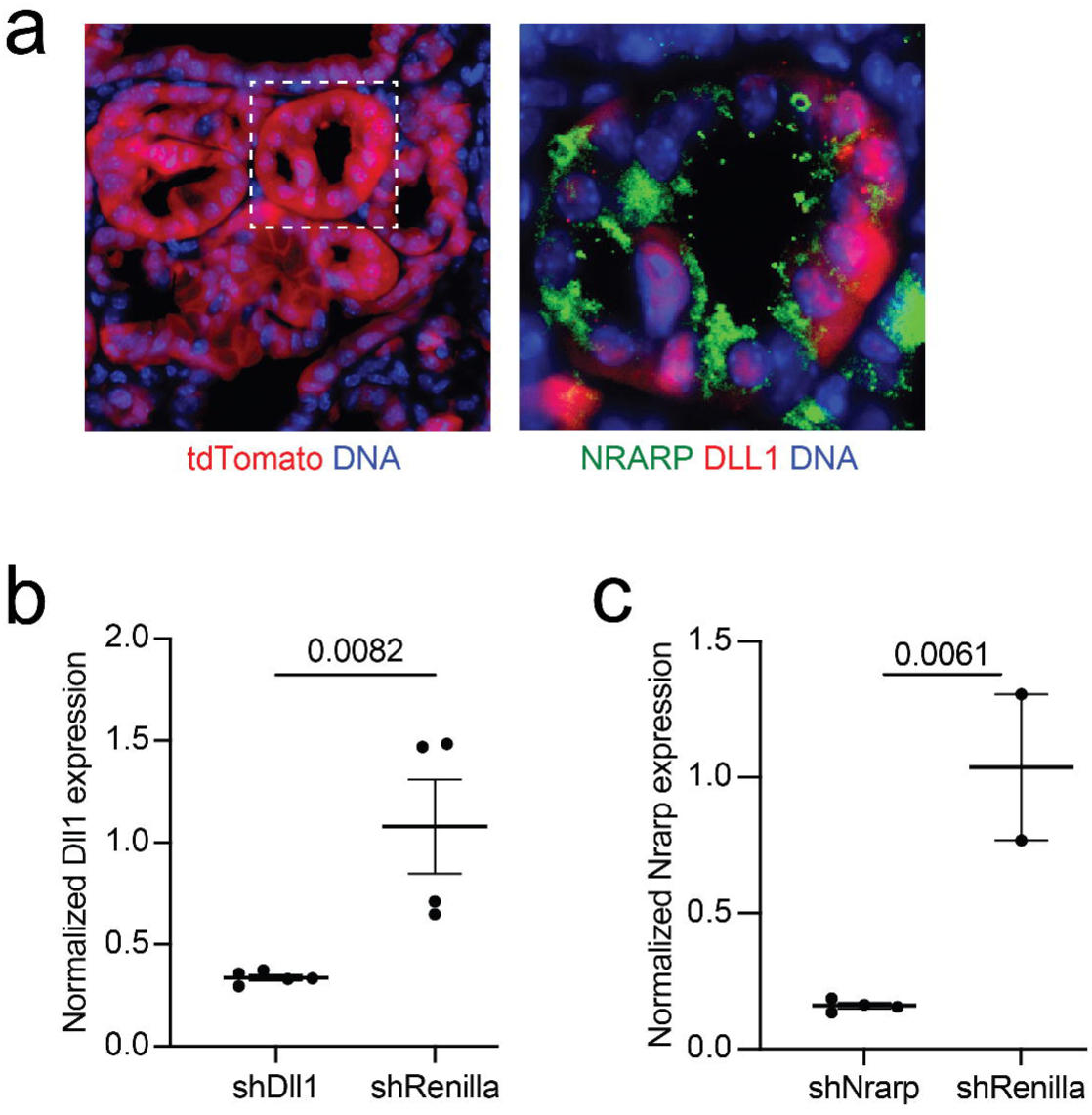
DLL1-Notch-NRARP signaling interactions between malignant cell states in PDAC, related to Figure 4. (**a**) Immunostaining for tdTomato (red) to detect cancer cells in an autochthonous *KPCT* PDAC tumor (left), inset: NRARP (green) and DLL1 (red) immunofluorescence. Scale bar: 50 µm. (**b**, **c**) *Dll1* (**b**) and *Nrarp* (**c**) expression assessed by quantitative PCR (qPCR) in orthotopic *7TCF/Porcn-DMA* reporter PDAC allografts harboring shDll1 (**b**) or shNrarp (**c**). Unpaired *t* tests were used to test significance in (**b**) and (**c**). Error bars indicate SEM.

**Figure S6.**
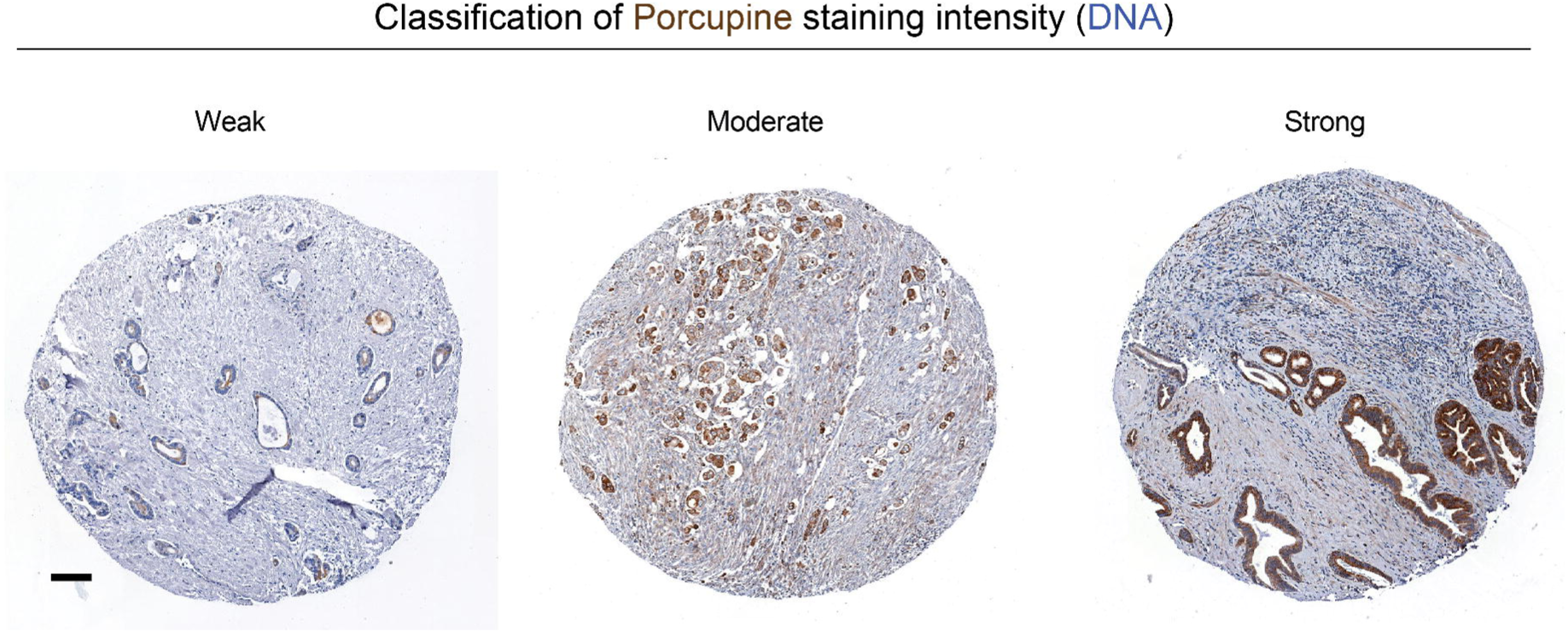
Induction of porcupine in a subset of cancer cells in human pancreas cancer. Representative immunohistochemical staining for porcupine (brown) in core biopsy samples in a human PDAC tissue microarray. Scale bar: 100 µm.

## Model 1. WNT-Sender cells grow as units emerging from proliferation of a Receiver cell

The dynamics of the Sender cell population:

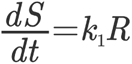

Here, the number of the Sender cells is assumed to grow in proportion to the number of the Receiver cells, with the coefficient *k*_1_.

The number of Receiver cells is assumed to grow in proportion to the number of competent Sender cells, and to decrease due to inter-conversion into the Sender cells. This latter process does not appreciably change the number of the Sender cells, and thus is not taken into the account in the first equation.

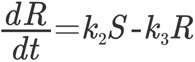

Assuming that the Receiver cells emerge and covert back to the Sender cells fast relatively to the growth of the Sender cells, we have approximately from the second equation:

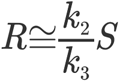

This relationship defines the relationships of the ratio of the Sender and Receiver cells.

The rate of the Sender cell multiplication is then given by the following equation:

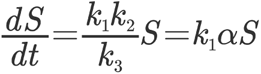

where *alpha* is the ratio of the Receiver and the Sender cells.

The number of the Sender cells then changes as:

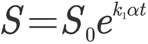

One can see that the growth is exponential, with the exponent proportional to the ratio of the Receiver to the Sender cells. Increasing this ratio artificially may increase the growth rate.

## Model 2. WNT-Sender cells grow independently of the Receiver cells

In this model, the Sender cells can divide independently of the Receiver cells. This changes the first equation of the above model to the following one.

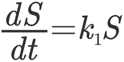

The second equation does not change, yielding the same and constant proportion of the Reviver to the Sender cell. However, the rate of the growth of the Sender cells no longer depends on the Receiver cells. This yields the following expression for the dependence of the number of the Sender cells on time:

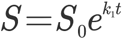

The relevance of the Receiver cells in this model is not clear.

## Notes

### Competing Interest Statement

The authors have declared no competing interest.

